# Transcriptomic mapping of the inter-individual variability of cellular stress response activation in primary human hepatocytes

**DOI:** 10.1101/2021.08.26.457742

**Authors:** Marije Niemeijer, Witold Wiecek, Suzanna Huppelschoten, Peter Bouwman, Audrey Baze, Céline Parmentier, Lysiane Richert, Richard S. Paules, Frederic Y. Bois, Bob van de Water

## Abstract

**Background & Aims:** One of the early key events of drug-induced liver injury (DILI) is the activation of adaptive stress responses, a cellular mechanism to overcome stress. Given the diversity of DILI outcomes and lack in understanding of population variability, we mapped the inter-individual variability in stress response activation to improve DILI prediction.

**Approach & Results:** High-throughput transcriptome analysis of over 8,000 samples was performed in primary human hepatocytes of 50 individuals upon 8 to 24 h exposure to broad concentration ranges of stress inducers: tunicamycin to induce the unfolded protein response (UPR), diethyl maleate for the oxidative stress response, cisplatin for the DNA damage response and TNFα for NF-κB signalling. This allowed investigation of the inter-individual variability in concentration-dependent stress response activation, where the average of benchmark concentrations (BMCs) had a maximum difference of 864, 13, 13 and 259-fold between different hepatocytes for UPR, oxidative stress, DNA damage and NF-κB signalling-related genes, respectively. Hepatocytes from patients with liver disease resulted in less stress response activation. Using a population mixed-effect framework, the distribution of BMCs and maximum fold change were modelled, allowing simulation of smaller or larger PHH panel sizes. Small panel sizes systematically under-estimated the variance and resulted in low probabilities in estimating the correct variance for the human population. Moreover, estimated toxicodynamic variability factors were up to 2-fold higher than the standard uncertainty factor of 10^1/2^ to account for population variability during risk assessment, exemplifying the need of data-driven variability factors.

**Conclusions:** Overall, by combining high-throughput transcriptome analysis and population modelling, improved understanding of variability in stress response activation across the human population could be established, thereby contributing towards improved prediction of DILI.

---

The development of drug-induced liver injury (DILI) is one of the main reasons for drug withdrawal from the market (1). Therefore, it is key to improve prediction of DILI at an early stage during drug development. For the evaluation of hepatotoxicity during the preclinical phase, primary human hepatocytes (PHHs) are currently considered to be the gold standard of tissue culture models for human liver toxicity. Despite their disadvantages, such as dedifferentiation and source limitations, they closely resemble human hepatocytes *in vivo*, allowing for the study of drug-induced hepatotoxicity (2).

Because of functional variability, often PHHs from three different individuals are used. However, the development of DILI can be highly patient specific and the inter-individual variability in DILI susceptibility is poorly understood. To account for this uncertainty, often a default uncertainty factor (UF) of 10 is used, with the goal to equally cover toxicokinetic and toxicodynamic variance (3). Whether this UF of 10 is enough to capture the full human population variance in hepatocyte function is unclear and this hampers a reliable DILI risk assessment. Therefore, there is a need for data-driven UFs which are both chemical and endpoint specific and accurately account for inter-individual variability.

During chemical-induced stress, cellular defensive mechanisms are activated to restore homeostasis. Monitoring the activation of these adaptive stress responses upon chemical exposure can give insight into DILI liabilities of chemical compounds as well as the underlying modes-of-action. Because of the protective functions of adaptive stress responses, inter-individual variations in their activities affect adverse outcomes such as DILI. Quantitative insight in these variations among the human population would allow the derivation of data-driven toxicodynamic UFs specifically for each type of stress response, thereby enabling better predictions of DILI liabilities.

Omic approaches such as transcriptomics, are powerful tools to fully map differences in adaptive stress response signaling networks across different patients. Several studies have already used transcriptomics approaches to compare differences between chemical-induced stress responses in different liver cell culture models (4,5). Due to advances in this area, novel approaches such as targeted TempO-seq technology can now be used, which allow for transcriptome mapping of gene sets of interest in a high-throughput fashion (6,7). This allows large-scale population studies and accurate analyses of inter-individual variance in transcriptomic perturbations for the improvement of DILI liability assessment. When combining experimental transcriptomic data with population modeling, an estimate of the variance across the entire population can be derived, an approach taken by Blanchette *et al*. for the evaluation of the variance in chemical-induced cardiotoxicity (8,9).

To map the inter-individual variability in stress response activation upon chemical exposure, we profiled the transcriptome for over 8,000 samples of a large panel of 50 cryo-preserved PHHs derived from different individuals and exposed to a broad concentration range of specific stress response inducing compounds. In combination with population modeling, these data allowed us to evaluate the influence of PHH panel sizes on the correct estimation of inter-individual variance in chemical stress responses and the need of data-driven toxicodynamic UFs, which will contribute to improved prediction of DILI liabilities.

## Experimental Procedures

### Cell culture

Plateable cryopreserved PHHs were derived from 54 different individuals (KaLy-Cell, Plobsheim, France) with permission of the national ethics committees and regulatory authorities (Supporting Data S1, Supporting Fig. S1). PHHs showing less than 70% confluency 24 h after plating were discarded for further analysis leading to a panel of PHHs derived from in total 50 individuals. Samples were also generated from PHHs directly upon thawing or from snap-frozen liver tissue derived from 8 individuals (Supporting Data S1).

### Cell treatment

After 24 h of plating, PHHs were exposed to four reference compounds or six DILI compounds, respectively (Supporting Table S1): tunicamycin, diethyl maleate (Sigma), TNFα (R&D systems), cisplatin (Ebewe), acetaminophen, propylthiouracyl, nitrofurantoin, ticlopidine, nefazodone and diclofenac (Sigma). To evaluate cytotoxicity, LDH release was evaluated in supernatants using a cytotoxicity detection kit (Roche).

### Transcriptomics dose-response analysis

The transcriptome was analyzed using the targeted TempO-seq technology (BioSpyder Technologies, Inc., Carlsbad, CA, USA) (6) with the supplemented S1500+ gene set of NIEHS (10) or targeted whole transcriptome panel (Supporting Data S2). Reads were aligned using the TempO-seq R package, normalized using the DESeq2 R package (11) and samples were filtered using a library size cut-off of 100.000 counts (Supporting Fig. S2). Benchmark concentration (BMC) modelling was done using the BMDExpress 2 software from Sciome LCC, NIEHS/NTP/NIH and EPA (12,13).

### Population modelling and predictive simulations

The BMC-maxFC values distributions were modelled in a Bayesian hierarchical framework. Large sample reference median coefficient of variation (CVs) were obtained for BMC and maxFC values by simulating of 1000 assays with hepatocytes from 2000 individuals each. For each simulated individual, BMC and maxFC values were simulated by Monte Carlo sampling using the average posterior estimates of the population parameters obtained by calibration of the model with the experimental data on 50 individuals. Similar assay simulations were performed for smaller, realistic, donor panel sizes (N = 3 to 50). CVs for BMC and maxFC values were obtained for each case.

Further details of methods can be found in supporting materials.

## Results

### Variability in basal gene expression levels across large panel of PHHs

As an initial step, the variance in dedifferentiation of PHHs derived from different individuals during culture was evaluated (Fig. 1A-B). For this purpose, we used whole transcriptome targeted TempO-seq analysis of liver tissue and derived cryopreserved PHHs upon thawing or cultured in 2D for 24 h for eight different individuals. Principal component analysis (PCA) showed greatest variability between individuals for freshly thawed PHHs (Fig. 1A-B). Noticeably, the transcriptome of PHHs cultured in 2D was more alike liver tissue compared to freshly thawed PHHs for principal component 1, representing 60.9% of the total variance (Fig. 1A). However, when only considering liver-related genes, both PHHs cultured in 2D or freshly thawed were similarly distinct from liver tissue for both principal components, although the latter being more variable (Fig. 1B). Genes involved in metabolism, such as *CYP3A4, CYP2C8* and *UGT2B7* were mostly differently expressed between liver tissue and PHHs, either cultured in 2D or freshly thawed.

**FIG. 1.**
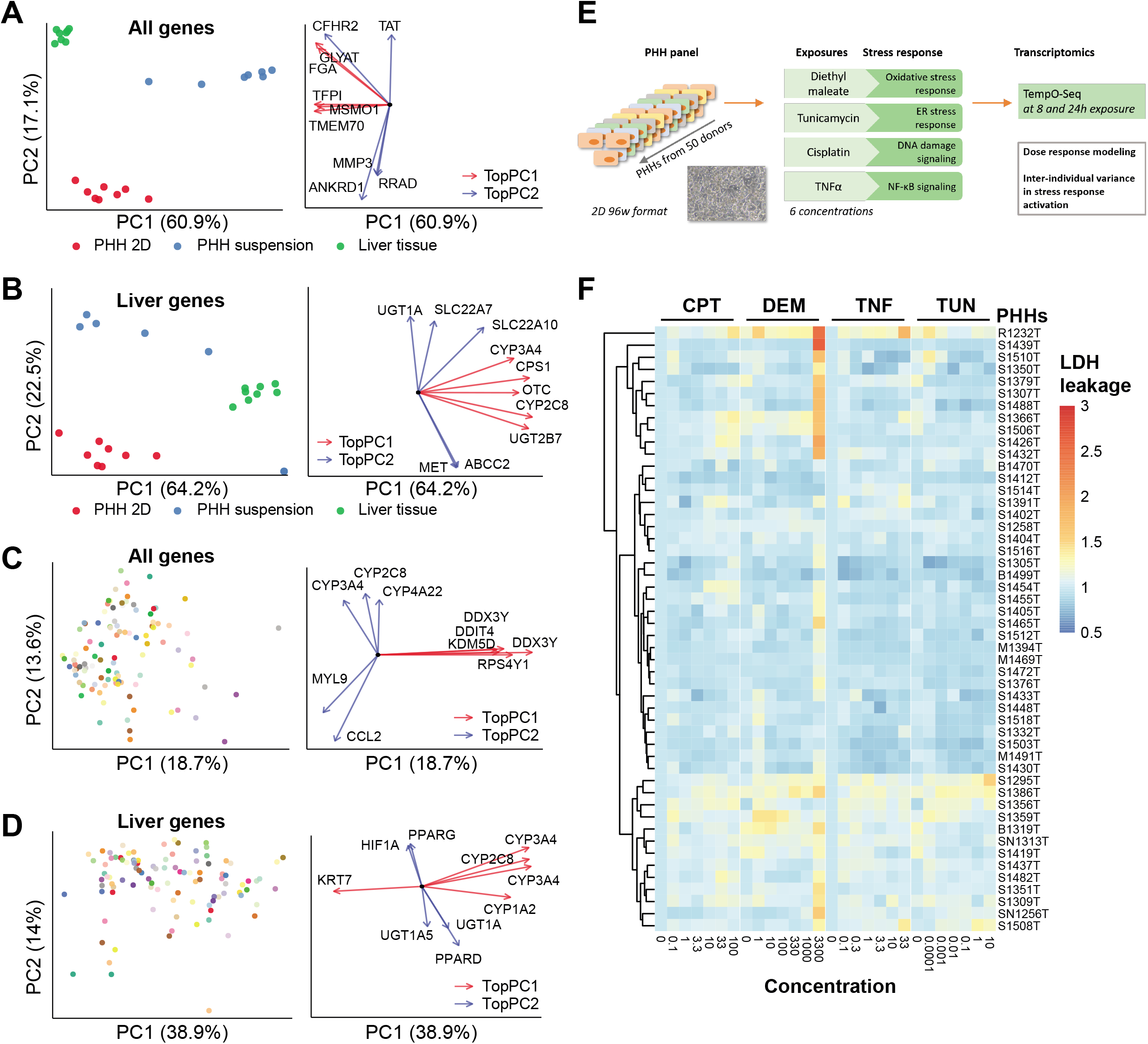
Characterization of the inter-individual variability utilizing a large panel of primary human hepatocytes derived from 50 individuals. A-D) Left panel: Principle component analysis (PCA) based on gene expression as log2 normalized counts. Right panel: Top 5 genes mostly determining PC1 or 2 depicted as vectors representing contribution and PC orientation. A-B) PCA of liver, PHHs in suspension or grown as 2D based on all genes (A) or liver-related genes (B) in whole transcriptome panel. C-D) PCA of panel of 50 PHHs depicted in different colours based on all genes (C) or liver-related genes (D) of S1500+ gene set. E) Schematic representation of experimental setup. F) LDH leakage upon exposure to DEM, TUN, CPT and TNFα in wide concentration range for 24 h across panel of PHHs. N = 3.

Next, we evaluated differences in gene expression for a panel of PHHs derived from 50 individuals cultured for 24 h in 2D for all genes (Fig. 1C), liver-related genes (Fig. 1D) or stress response-related genes (Supporting Fig. S3). PHHs from some individuals showed altered expression for genes dominating the variance across the first principal component based on all genes, such as *DDIT4*, a regulator of mTOR activity induced upon various stress conditions (Fig. 1C). In addition, variability in various phase-I enzymes such as *CYP3A4* was seen across the panel of PHHs mostly affecting the second principal component for liver-related genes (Fig. 1D and Supporting Fig. S4). Also, large variation was seen in the expression of *CCL2*, a chemokine involved in the recruitment of monocytes and basophils, affecting the principal components for all genes (Fig. 1C) but also more specifically UPR and NF-κB signaling related genes (Supporting Fig. S3).

### Difference in sensitivity towards chemical-induced cell death

To evaluate the inter-individual variability in chemical-induced cellular stress responses across the panel of 50 PHH cultures, PHHs were exposed for 8 or 24 h to broad concentration ranges of specific stress inducers: diethyl maleate (DEM) to induce the oxidative stress response, tunicamycin (TUN) for the unfolded protein response, cisplatin (CPT) for the DNA damage response and TNFα for inflammatory NF-κB signalling (Fig. 1E). Great difference in viability, measured by LDH release, was only seen between PHHs from different individuals following 24 h exposure at the highest concentration of DEM, where significant induction of cell death was seen for a subset of PHHs while other PHHs did not show viability loss at all (Fig. 1F). The other compounds did not lead to significant loss of viability upon 24 h of exposure, although some PHHs were more sensitive, such as R1232T and S1295T, resulting in a minor LDH release at the highest concentrations.

### Inter-individual variability in chemical-induced stress response activation

Next, the effects of 8 or 24 h of chemical exposure on the PHH transcriptomes were analysed. Exposure to the oxidative stress-inducing compound DEM resulted in clear upregulation of NRF2 target genes *HMOX1* and *SRXN1* at concentrations ranging from 330 μM to 3300 μM for both time points (Fig. 2A). Cisplatin-induced DNA damage signalling in the panel of PHHs was most profound at a concentration of 10 μM upon 24 h of exposure, where in general the highest upregulation of P53 target genes *BTG2* and *MDM2* was seen. PHHs showed variable responses, where some showed *BTG2* or *MDM2* upregulation already at a cisplatin concentration of 1 μM, while others required higher amounts of this compound. Regarding the UPR induced by TUN, activation of both the adaptive gene *HSPA5* and pro-apoptotic related gene *DDIT3* was seen at a concentration of 0.01 μM or higher at the 24 h time point. Some PHHs showed already strong upregulation of both genes at 0.01 μM of TUN, while other PHHs only showed upregulation at 1 μM or higher, suggesting particularly large differences in the UPR response of different individuals. Upon TNFα exposure to study variance in inflammation signalling, NF-κB target genes *ICAM1* and *TNFAIP3* were activated in a concentrationdependent manner. For most of the PHHs, activation was seen at 1 ng/mL and higher when exposed for 24 h. Some PHHs showed activation at the lowest concentration of 0.1 ng/mL, while other PHHs were completely unresponsive to TNFα exposure.

**FIG. 2.**
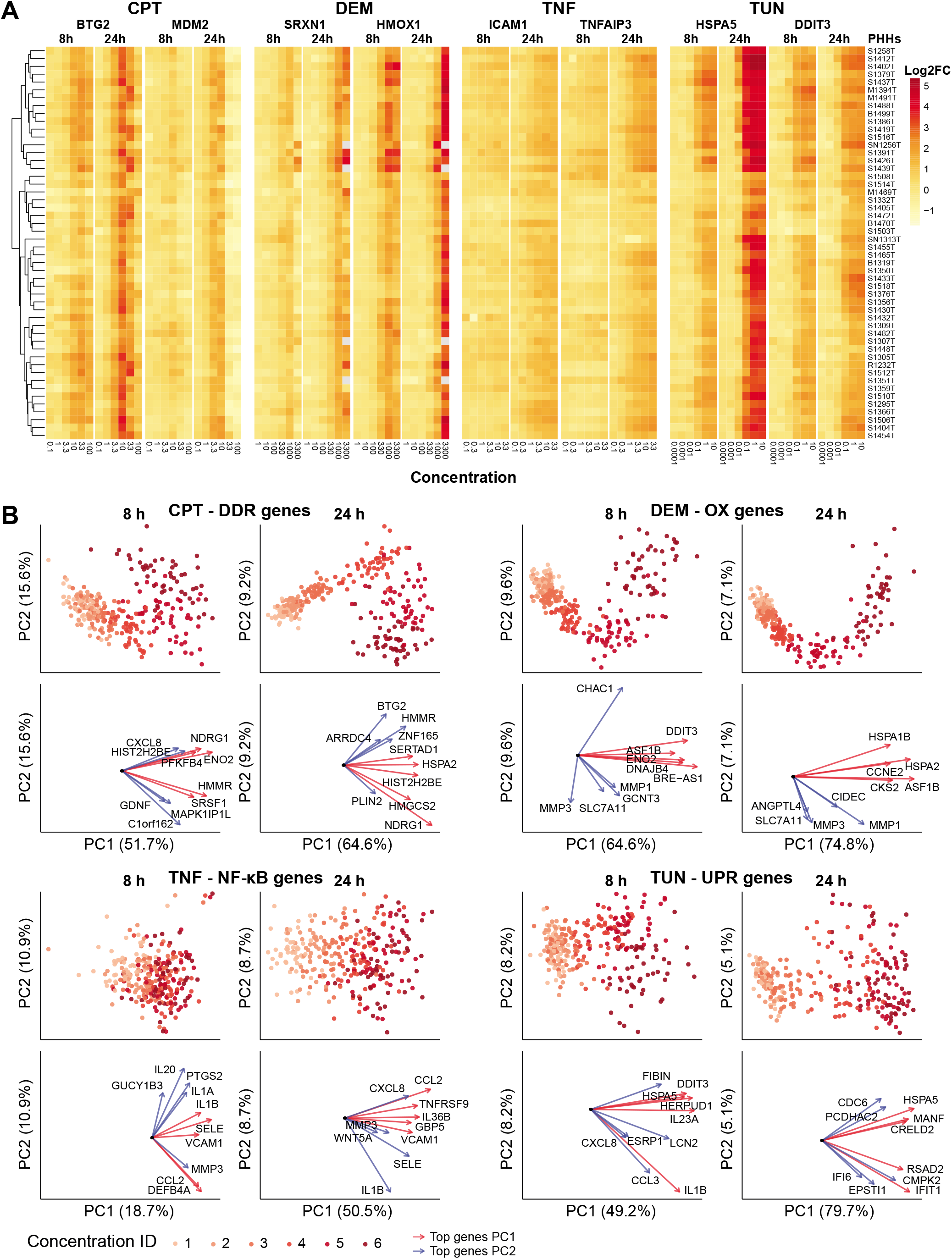
Inter-individual variability in concentration-dependent stress response activation. A) Hierarchical clustering of log2 fold change gene expression of key genes for each evaluated stress response pathway of panel of PHHs exposed to reference compounds. B) Principle component analysis (PCA) based on log2 fold changes of stress responsive genes of panel of PHHs exposed to DEM, TUN, CPT and TNFα. Top 5 genes mostly determining PC1 or 2 depicted as vectors representing contribution and PC orientation. N=3.

Subsequently, we selected the top 50 most strongly activated genes at the lowest effective concentration (Supporting Data S2) of each stress response reference compound. The inter-individual variability in the concentration-dependent activation of these top 50 genes for each stress response was evaluated by PCA (Fig. 2B). In general, for all compounds a clear concentration-dependent shift was seen for all PHHs. Most variability was observed between different PHHs for TUN-induced UPR genes and TNFα-induced NF-κB related genes, mainly driven by variable expression of cyto- or chemokines such as *IL1B*.

### Inter-individual differences in points-of-departure of chemical-induced stress response activation

To evaluate the inter-individual variability in sensitivity to chemical-induced stress response activation, we defined benchmark concentrations (BMCs) at which the top 50 most strongly activated genes for each specific compound showed an increase of 1 standard deviation in gene expression (Supporting Data S3). Upon dose response modelling, we observed large differences in the gene-specific BMCs between different PHHs (Fig. 3A). In general, most variability was seen at 8 h of exposure, while at 24 h the response was more stable across the PHH panel. Large shifts in the BMC distribution could be seen especially at the 8 h time point, with medians varying 864, 259, 13 and 13-fold for TUN, TNFα, DEM and CPT, respectively (Table 1). This means that BMC estimations can shift significantly depending on which PHHs are taken along in chemical toxicity testing at early time points. Variability could also be observed in the distribution of the maxFC of the top 50 most strongly activated genes among the different PHHs, where compound sensitivity correlated positively with levels of upregulation (Supporting Fig. S6).

**FIG. 3.**
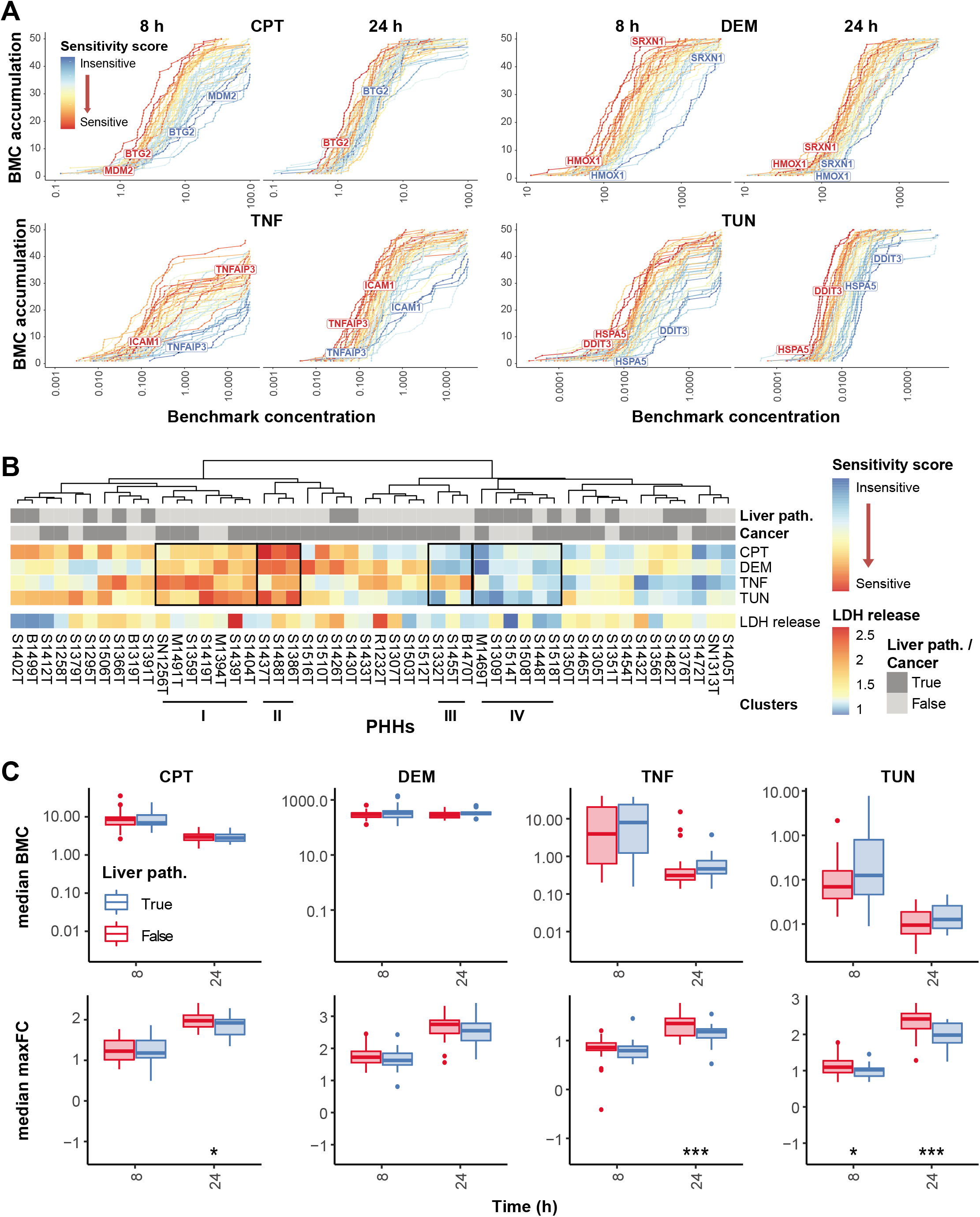
Large shift in bench mark concentration distribution across panel of primary human hepatocytes (PHHs) affected by liver disease status. A) Distribution of bench mark concentrations (BMC) of top 50 stress responsive genes for each compound. Lines represent the different PHHs within panel. B) Hierarchical clustering of sensitivity score for panel of PHHs for each treatment. The LDH leakage is shown of 3300 μM DEM at 24 h and the disease status. C) Boxplots of median BMC or maxFC of top 50 stress responsive genes of exposure for each treatment for PHHs with or without liver disease status. Significance levels represented as * p < 0.1, ** p < 0.05, *** p < 0.01. N = 3.

**Table 1.** Median benchmark concentrations for each PHH based on top 50 stress responsive genes for each compound and time point.

|  | *CMP* | *Median BMC ± SD*<br>*(min-max)* | *FC min-max*<br>*Median BMC* | *TDVF<sub>0.01</sub>*<br>*Exp data* | *TDVF<sub>0.01</sub>*<br>*Model\** | *SD*<br>*TDVF<sub>0.01</sub>*<br>*Model\** |
| --- | --- | --- | --- | --- | --- | --- |
| **8 h** | TUN | 0.534 ± 1.320<br>(0.009 - 7.768) | 864.1 | **6.545** | **6.317** | 0.582 |
|  | DEM | 352.486 ± 243.643<br>(111.726 - 1405.220) | 12.6 | 2.517 | 1.828 | 0.066 |
|  | CPT | 9.488 ± 5.867<br>(2.640-35.447) | 13.4 | 2.786 | **3.241** | 0.202 |
|  | TNF | 12.447 ± 13.310<br>(0.157 - 40.633) | 258.8 | **26.524** | **5.315** | 0.423 |
| **24 h** | TUN | 0.023 ± 0.061<br>(0.002 - 0.447) | 223.5 | **4.950** | **4.811** | 0.386 |
|  | DEM | 315.597 ± 108.883<br>(176.656 - 623.096) | 3.5 | 1.701 | 1.624 | 0.047 |
|  | CPT | 3.083 ± 1.022<br>(1.471 - 6.401) | 4.4 | 1.847 | 2.184 | 0.090 |
|  | TNF | 0.926 ± 2.252<br>(0.137 - 15.263) | 111.4 | 2.516 | 2.556 | 0.128 |

*\*Based on population modelling of 2000 virtual donors for 1000 assays*

To further assess differences in PHH sensitivity, sensitivity scores were determined based on both the median BMC and maxFC for the top 50 activated genes for each compound for both time points (Supporting Fig. S5B and Data S4). A subset of PHHs (cluster I-II) showed high sensitivity scores for all stress responses induced by the reference compounds and also showed higher cell death induction at the highest concentration of DEM (Fig. 3B). In addition, a subpanel of PHHs (cluster III) was highly sensitive towards TNFα, while being insensitive towards all other chemical induced stress responses. Thus, PHHs can be sensitive to chemical compounds in general, but also selectively sensitive for specific types of stress-inducing chemicals. In addition, negative correlations could be seen between cell death induction as measured by LDH release and the sensitivity scores, where in general PHHs with high LDH release were more sensitive for stress response activation having lower sensitivity scores (Fig. 3B, Supporting Fig. S7). This indicates that for the most sensitive PHHs the switch towards adverse signalling was already reached at lower concentrations of stress-inducing chemicals.

Next, we calculated a data-driven toxicodynamic variability factor (TDVF_0.01_), which is the ratio between the median population point-of-departure (PoD) and the 1% quantile individual PoD. This TDVF_0.01_ accounts for underestimation of the variance within the human population when estimating the PoDs for cellular stress response activation upon chemical exposures (Table 1). For DEM-mediated oxidative stress and cisplatin-induced DNA damage, the commonly used uncertainty factor of 3.16 (3) would be enough since data-based TDVF_0.01_s were all lower at both 8 and 24 h timepoint. However, for 8 h TUN-induced UPR activation and TNFα-mediated NF-κB signaling, the defined TDVF_0.01_ was higher than the uncertainty factor of 3.16, namely 6.5 and 26.5, respectively. At the 24 h timepoint, which showed in general less variability across the panel of PHHs than the 8 h timepoint, TUN-induced UPR had a TDVF_0.01_ of 5, which is also higher than the standard factor. These examples show that the standard uncertainty factor is not enough to capture the variation in all chemical-induced stress responses within the human population.

### Influence of pathological background on sensitivity of chemical-induced stress response activation

Next, the influence of the disease status, such as cancer or different types of liver pathology, on stress response activation upon chemical exposure in PHHs was evaluated. In general, the presence of any type of cancer did not have an obvious influence on the sensitivity of the PHHs towards chemical-induced stress, since clear clustering of sensitivity scores of PHHs derived from patients with cancer was not seen (Fig. 3B). In addition, presence of cancer only mildly influenced the distribution of median maxFC and BMC for the top 50 stress responsive genes (Supporting Fig. S8B). In contrast, PHHs from patients having any type of liver pathology were in general less sensitive towards chemical-induced stress, especially for TUN-induced UPR and TNFα-induced NF-κB signalling (Fig. 3B). When we evaluated the distribution of the median maxFC, a significant difference could be seen between PHHs derived from patients with or without liver pathology when exposed to TUN or TNFα for 24 h (Fig. 3C). The same trend was seen for DEM and CPT, although this was not significant. Possibly, PHHs from patients with a certain liver pathology were already at a higher level of stress leading to lower fold changes upon chemical-induced stress. Indeed, these PHHs showed higher basal expression of the top 50 stress responsive genes compared to PHHs without liver pathology (Supporting Fig. S8C). Variability in differentiation status did not have an effect on BMC and maxFC distribution (Supporting Fig. S9).

### Variance in pathway enrichment upon chemical exposure across panel of PHHs

To get more insight in chemical-induced stress pathway activation, gene set enrichment analysis of gene ontology terms was performed using the maxFC as input for ranking of the genes. In general, upon exposure to each reference compound, expected terms were enriched related to anticipated chemical-induced stress response pathways (Fig. 4A, Supporting Fig. S10). For instance, TUN treatment led to the strong enrichment of ER stress and UPR-related terms, TNFα resulted in chemotaxis and chemokine-related terms, DEM gave enrichment of heat shock or ion response-related terms and CPT treatment led to enrichment of ion or metabolic-related terms. All chemicals led to enrichment for both sensitive and insensitive PHHs, although some terms were more specific for either sensitive or insensitive PHHs, e.g. terms related to response to ions were more enriched for insensitive PHHs upon DEM exposure (Fig. 4A). Clustering of enriched terms for at least three PHHs showed specific enrichment of terms related to ribosomal localization or subunits for insensitive PHHs upon TUN or CPT exposure, while terms related to response to virus or interferon were enriched for specifically sensitive PHHs upon TNFα treatment (Fig. 4B).

**FIG. 4.**
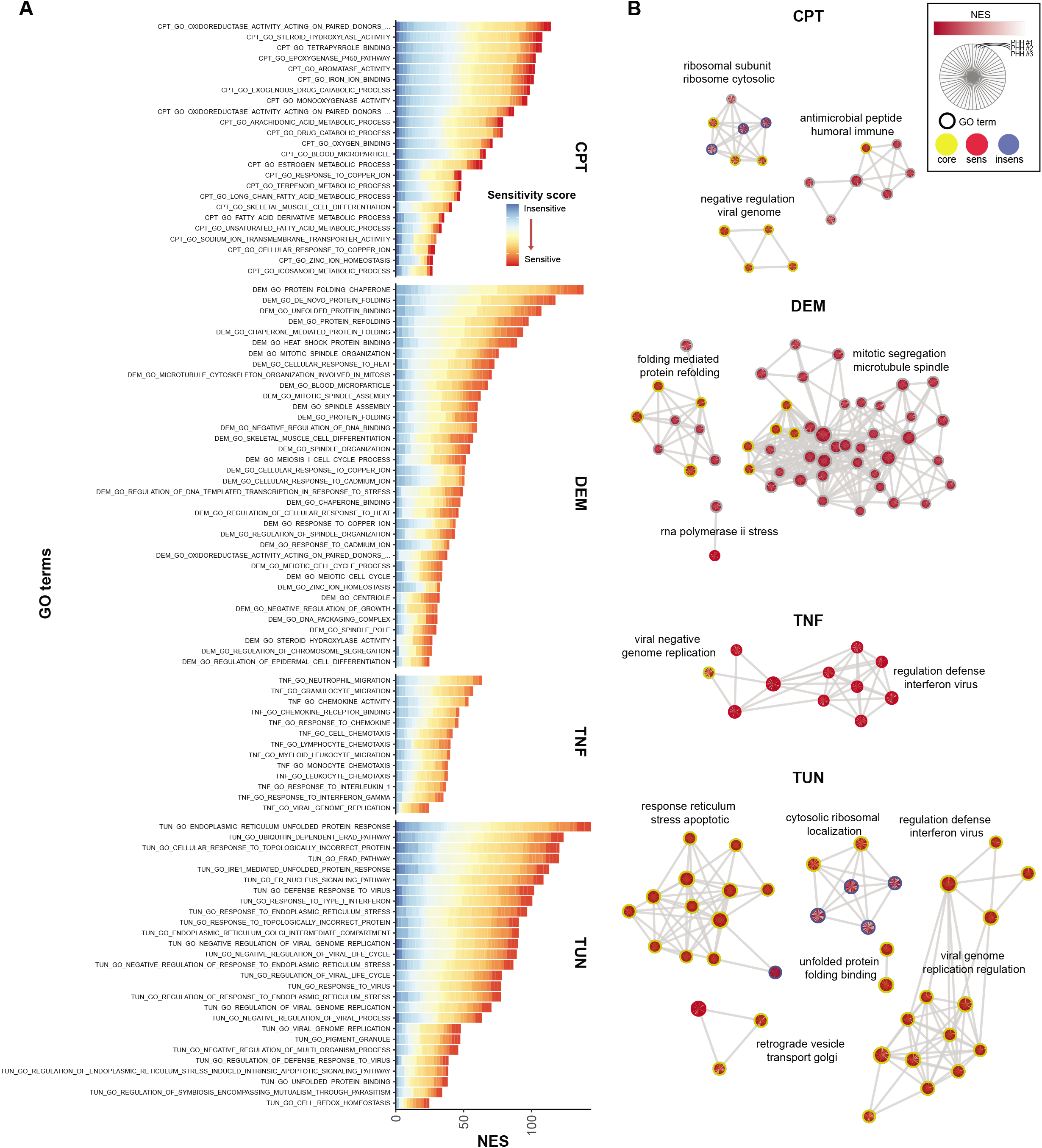
Variance in gene set enrichment upon chemical-induced stress. Gene set enrichment analysis (26) was done using gene ontology terms v7.1 and the maximal fold change across concentration range (maxFC) of measured S1500+ genes as input for each PHH and treatment for 24h with each reference compound. A) Barplots of normalized enrichment scores (NES) of significantly enriched GO terms with a cut-off of adjusted FDR of < 0.05 for at least 10 different PHHs. NES were stacked for all PHHs showing significantly enrichment for each term. B) Significantly enriched terms in which one or more terms within cluster showed specific enrichment in either most sensitive (red circle) or insensitive (blue circle) PHHs, or both defined as core (yellow circle) using Cytoscape (27) and EnrichmentMap (28). Within each term, NES for each PHH is depicted in grey to red scale from low to high. Enriched terms were clustered and summarized with 3 to 4 keywords using WordCloud (29).

### Inter-individual variability in stress response activation by hepatotoxicants

The inter-individual difference in sensitivity towards chemical-induced oxidative stress response and UPR activation was further analyzed by screening the three most sensitive and insensitive PHHs with various hepatotoxicants known to induce oxidative stress (acetaminophen, propylthiouracyl and nitrofurantoin) or ER stress (ticlopidine, nefazodone and diclofenac) (Fig. 5). Upon evaluation of differences in cell death induction, a concentration-dependent increase in LDH release was seen for all tested hepatotoxicants, except for acetaminophen and propylthiouracil (Fig. 5A). Both nitrofurantoin and nefazodone already showed cell death induction at 25× Cmax or higher. The most sensitive PHHs showed higher cytotoxicity when exposed to nitrofurantoin and diclofenac than the most insensitive PHHs. For the other hepatotoxicants, no difference was seen. Next, the expression of the top 50 activated genes was evaluated for both the oxidative stress response and UPR, where in particular nitrofurantoin showed higher induction in sensitive PHHs compared to insensitive (Fig. 5B). Other hepatotoxicants did not show clear distinction when only evaluating these genes. Thereafter, the difference in the distribution of BMCs and maxFC of all responsive genes was evaluated between the most sensitive and insensitive PHHs for each tested DILI compound. The hepatotoxicants propylthiouracyl, acetaminophen and ticlopedine showed clear separation for most of the sensitive and insensitive PHHs, but not all, in BMC and maxFC distribution (Fig. 5C, Table 2). For all other hepatotoxicants, differences in BMCs and maxFCs were more variable.

**FIG. 5.**
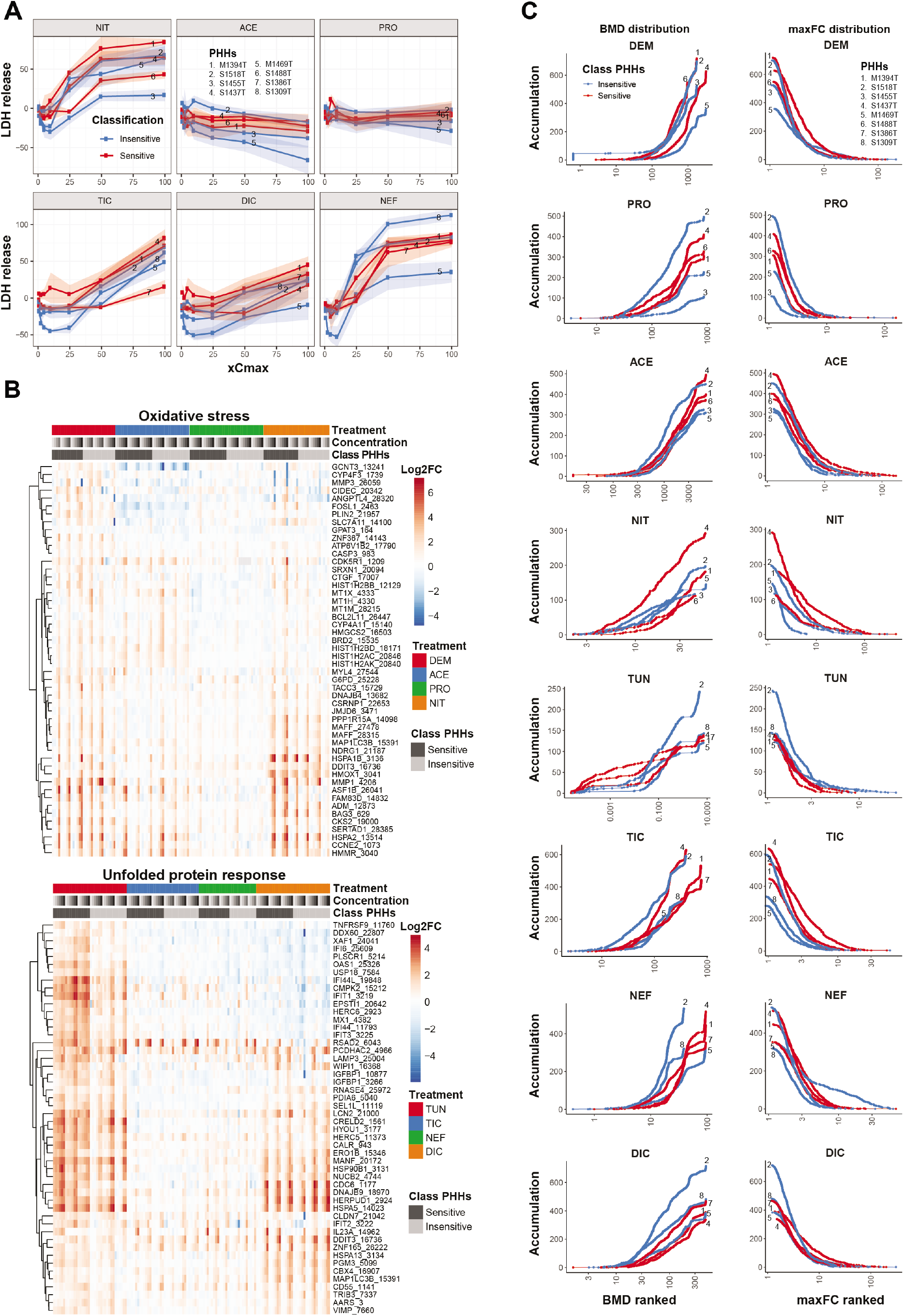
Difference in sensitivity towards hepatotoxicants within a subpanel of primary human hepatocytes (PHHs). A) LDH release upon treatment with hepatotoxicants for 24 h in 6 defined most sensitive or insensitive PHHs. B) Hierarchical clustering of log2 fold changes of top 50 stress responsive genes for the oxidative stress and unfolded protein of PHHs exposed to various hepatotoxicants (acetaminophen; ACE, propylthiouracyl; PRO, nitrofurantoin; NIT, ticlopidine; TIC, nefazodone; NEF, diclofenac; DIC) for 24 h. C) Distribution of benchmark concentration (BMC) and maximal fold change across concentration range (maxFC) for all responsive genes of PHHs exposed to various hepatotoxicants for 24 h. N = 3.

**Table 2.** Median maximal fold change across concentration range (maxFC) and benchmark concentration (BMC) for most sensitive and insensitive PHHs based on all responsive genes for each treatment.

| *CMP* | *Median maxFC* |  |  |  |  |  | *Median BMC (μM)* |  |  |  |  |  |
| --- | --- | --- | --- | --- | --- | --- | --- | --- | --- | --- | --- | --- |
|  | *Insensitive PHHs* |  |  | *Sensitive PHHs* |  |  | *Insensitive PHHs* |  |  | *Sensitive PHHs* |  |  |
| **OX** | *M1469T* | *S1455T* | *S1518T* | *M1394T* | *S1437T* | *S1488T* | *M1469T* | *S1455T* | *S1518T* | *M1394T* | *S1437T* | *S1488T* |
| DEM | 2.94 | 2.51 | 2.25 | 2.33 | 3.48 | 2.53 | 1024.47 | 580.07 | 629.48 | 672.95 | 1051.43 | 375.30 |
| ACE | 3.14 | 2.95 | 3.59 | 2.93 | 3.61 | 3.68 | 1528.25 | 1331.12 | 1069.92 | 1761.77 | 1993.41 | 1379.58 |
| NIT | 1.92 | 3.10 | 3.17 | 6.31 | 2.86 | 3.80 | 20.65 | 11.60 | 24.03 | 29.02 | 17.37 | 22.12 |
| PRO | 1.96 | 1.54 | 2.16 | 1.78 | 2.03 | 2.22 | 293.33 | 345.93 | 193.57 | 312.23 | 311.24 | 235.90 |
| **UPR** | *M1469T* | *S1309T* | *S1518T* | *M1394T* | *S1386T* | *S1437T* | *M1469T* | *S1309T* | *S1518T* | *M1394T* | *S1386T* | *S1437T* |
| TUN | 1.84 | 1.98 | 1.78 | 1.73 | 1.89 | 1.78 | 0.07 | 0.21 | 0.19 | 0.004 | 0.19 | 0.04 |
| DIC | 3.08 | 2.68 | 2.88 | 5.02 | 2.90 | 3.70 | 122.25 | 72.45 | 68.45 | 137.28 | 99.12 | 137.80 |
| NEF | 3.33 | 2.13 | 2.09 | 2.92 | 2.61 | 2.42 | 37.31 | 18.24 | 19.01 | 37.25 | 25.66 | 40.28 |
| TIC | 1.97 | 2.20 | 2.21 | 3.63 | 2.67 | 2.48 | 103.17 | 141.57 | 96.00 | 229.69 | 210.65 | 151.71 |

*\*Diethyl maleate; DEM, acetaminophen; ACE, propylthiouracyl; PRO, nitrofurantoin; NIT, tunicamycin; TUN, ticlopidine; TIC, nefazodone; NEF, diclofenac; DIC.*

### Influence of PHH panel size on probability of estimating correct variance of stress response activation

To check the inter-individual variability model for the entire human population in chemical-induced stress response activation, both the BMC and maxFC for the top 50 stress responsive genes for each compound was simulated for 500 virtual PHHs. The distribution of the simulated BMCs and maxFCs were in concordance with the experimental data based on the panel of 50 PHHs showing similar distributions (Supporting Fig. S11-12).

To evaluate the effect of the number of PHHs from different individuals used to obtain the correct population CVs in stress response activation, reference (large sample size) population variances of BMC and maxFC values for the top 50 stress responsive genes were first estimated by simulating 1,000 assays with a panel of PHHs from 2,000 virtual individuals (Fig. 6A-B).

**FIG. 6.**
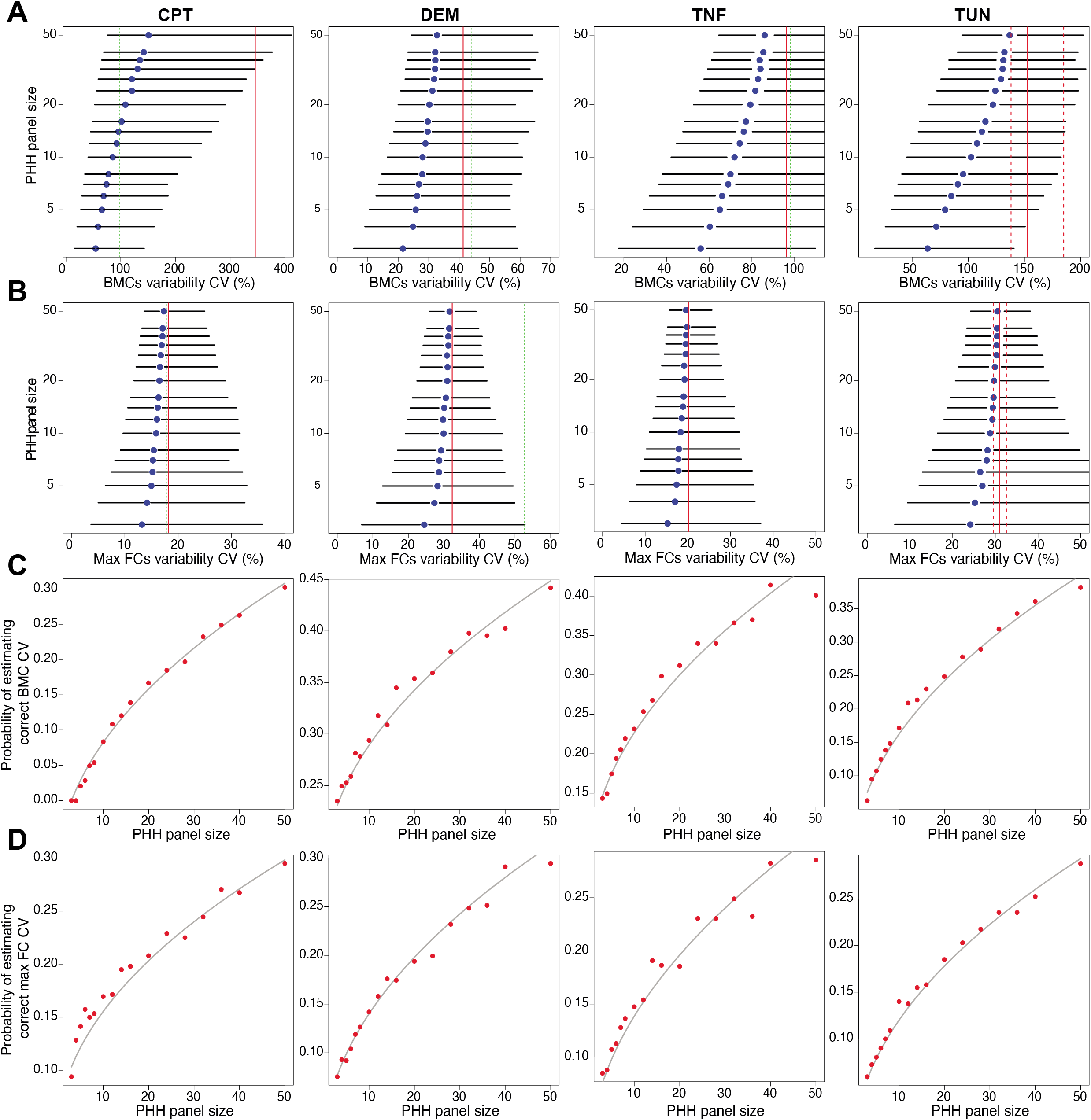
Influence of primary human hepatocyte (PHH) panel size on probability to capture correct variance in stress response activation. A-B) Forest plots of simulated distributions (medians ± 5^th^ / 95^th^ percentiles) of the estimated coefficients of variation (CVs) of benchmark concentrations (BMC) (A) and maximal fold changes across concentration range (maxFC) (B) based on top 50 stress responsive genes for each reference compound as a function of PHH panel size. As reference, 1000 assays with a panel of PHHs from 2000 individuals was simulated depicted as red lines (median: solid ± 5^th^ / 95^th^percentiles: dashed). C-D) Probability of reporting a correct CV falling between the 5^th^ and 95 ^th^ percentile of the reference CVs as a function of PHH panel size for BMC (C) and maxFC (D). Red dots resemble Monte Carlo simulation estimates and grey lines resemble visually fitted smoothing curves.

Next, inter-individual CV estimates were simulated when using different panel sizes of PHHs derived from 3 to 50 individuals for each reference compound (Fig. 6A-B). When the PHH panel size was increased, the CV estimates became increasingly precise and converge to the true human population variability. For small PHH panel sizes which are commonly used during hepatotoxicity testing, the imprecision is very large, leading to consistent underestimation of the inter-individual variance in stress response activation upon chemical exposure.

Next, the probability of obtaining a CV which is close to the true human population CV was evaluated when using different PHH panel sizes (Fig. 6C-D). Overall, when using a PHH panel size of 10 or less the probability to estimate the correct CV of the BMC is very low, namely smaller than < 0.3, 0.05, 0.15 and 0.2 for DEM, CPT, TUN and TNFα, respectively (Fig. 6C). When using a PHH panel size of 50, the maximal probability that could be reached to estimate the correct CV of the BMC is < 0.45, 0.3, 0.4 and 0.4 for DEM, CPT, TUN and TNFα, respectively, still resulting in quite some uncertainty. For estimating the correct CV of the maxFC, a maximal probability of 0.3 was reached for all compounds when using a PHH panel size of 50, which is also still quite low (Fig. 6D). Therefore, we propose to use an additional uncertainty factor to account for the uncertainty in estimating the true median population PoD and variance of stress response activation upon chemical exposure during hepatotoxicity testing in PHHs. Indeed, simulation of 2,000 virtual donors for 1,000 assays resulted in TDVF_0.01_s that were up to 2-fold higher than the standard used UF of 3.16 for TUN, CPT and TNFα for 8 h and TUN for 24 h (Table 1).

## Discussion

Human tissue-culture based methods hold great promise for the prediction of DILI liabilities at an early stage during drug development. Here, we evaluated the effects of inter-individual differences in sensitivity to chemical-induced stress in PHHs, to determine data-driven toxicodynamic uncertainty factors for specific types of DILI-relevant stress responses. In our analysis, we focussed on compounds inducing the UPR, the oxidative stress response, the DNA damage response and cytokine-mediated NF-κB signaling. The activation of each of these stress responses is considered a key event leading to the development of DILI (14). Moreover, the consequent upregulation of a specific set of gene transcripts can be used as a potential mode-of-action revealing endpoint for the evaluation of hepatoxicity (14,15). Our results indicate that large differences in the distribution of BMCs are observed between different PHHs. Moreover, the population modelling revealed large under-estimation of the inter-individual variance of stress response activation when small PHH panel sizes were used.

Our data exemplify the need of an accurately determined data-driven uncertainty factors during risk assessment to improve the prediction of DILI liabilities. Our results indicate that these uncertainty factors can be at least 2-fold higher than the standard toxicodynamic uncertainty factor of 10^1/2^. In concordance, Blanchette *et al*. characterized the human population variance of chemical-induced cardiotoxicity using hiPSC-derived cardiomyocyte panel together with population modelling and also showed higher TDVFs than the standard 10^1/2^ (8,9). Likewise, Abdo *et al*. found that some of the 179 screened chemicals led to TDVFs higher than 10 when evaluating cytotoxicity in 1,086 lymphoblastoid cell lines (16).

Potentially, several factors may influence PHH sensitivity to chemical-induced stress, including differences in health characteristics of PHH donors. We found that PHHs from patients with any type of liver disease showed less induction of stress responses upon exposure, especially for TNFα-induced NF-κB signalling and TUN-induced UPR. These PHHs showed already higher expression of stress response-related genes in control conditions, thereby leading to less capacity to further induce protective stress responses upon chemical exposure. Indeed, there are strong correlations between liver diseases, such as non-alcohol fatty liver disease, non-alcohol steatohepatitis and liver cancer, increased activation of e.g. the UPR and the inflammatory response (17–20), and increased DILI susceptibility (21,22). Thus, the liver disease background of PHHs can have a significant impact on their sensitivity to DILI compounds. In this respect, special attention should be given to the inflammatory state of the liver prior to the isolation of PHHs. Large differences were seen in basal expression of chemokines, such as *CCL2*, possibly affecting their sensitivity towards chemical exposure. Indeed, the presence of inflammation is one of the susceptibility factors for development of DILI (23,24). In addition, large variability was also seen in the expression of inflammatory genes upon chemical exposure across panel of PHHs. Inflammatory genes such as *IL1B* and *CCL3* were most differently expressed across panel of PHHs upon TUN treatment. Several studies have shown the relation between UPR activation and inflammatory signalling through activation of NF-κB and the NLRP3 inflammasome, leading to induction of *IL1B* expression (25).

In conclusion, we demonstrated that DILI associated stress response activation is highly variable between different individuals. This highlights the need to use toxicodynamic uncertainty factors for safety evaluation. We show that the currently used standard uncertainty factor of 10^1/2^ is not sufficient to capture all variance for every chemical or endpoint measured in PHHs. For activation of the UPR and NF-κB signalling the defined TDVF was up to 2-fold higher than this standard. This exemplifies a general need for the use of data-driven mechanism-specific uncertainty factors to accurately correct for toxicodynamic variance across the human population to improve assessment of DILI liabilities.

## List of Abbreviations

ACE: acetaminophen
BMC: benchmark concentration
CPT: cisplatin
CV: coefficient of variation
DEM: diethyl maleate
DIC: diclofenac
DILI: drug-induced liver injury
GSEA: gene set enrichment analysis
LDH: lactate dehydrogenase
maxFC: maximum fold change across concentration range
NEF: nefazodone
NIT: nitrofurantoin
PCA: principal component analysis
PHHs: primary human hepatocytes
PoD: point-of-departures
PRO: propylthiouracyl
TDVF: toxicodynamic variability factor
TIC: ticlopidine
TUN: tunicamycin
UF: uncertainty factor
UPR: unfolded protein response

