## Supplemental material for "Transcriptomic mapping of the inter-individual variability of cellular stress response activation in primary human hepatocytes"

### Supporting Information

#### Supporting Experimental Procedures

##### *Cell culture*

Plateable cryopreservered PHHs were derived from 54 different individuals provided by KaLy-Cell (Plobsheim, France) with permission of the national ethics committees and regulatory authorities (Supporting Data S1). For plating, PHHs were thawed in warm waterbath at 37°C and diluted in pre-warmed thawing UCRM medium (IVAL, Columbia, USA) followed by centrifugation at 170g for 20 min at RT. Thereafter, cell pellet was diluted in seeding UPCM medium (IVAL) and viability was assessed using the Trypan blue exclusion method. Cells were plated at a density of 70.000 cells in 100 µL per well in 96 wells BioCoat Collagen I Cellware plates from Corning (Wiesbaden, Germany). After 6 hours, medium was refreshed with seeding UPCM medium. PHHs showing less than 70% confluency 24 h after plating were discarded for further analysis leading to a panel of PHHs derived from in total 50 individuals. Characteristics of these PHHs are depicted in Supporting Fig. S1. To evaluate variability in dedifferentiation upon culturing of PHHs, samples were also generated from PHHs in suspension directly upon thawing or from snap-frozen liver tissue derived from 8 individuals (Supporting Data S1).

##### *Cell treatment*

Prior to compound exposure, PHHs were first washed after 24 h of attachment using 1x PBS to remove unattached cells. Exposures were done using William's E medium supplemented with 100 U/mL penicillin and 100 µg/mL streptomycin. PHHs were exposed to four reference compounds in a broad concentration range known to induce specific stress response pathways (Supporting Table S1), namely tunicamycin and diethyl maleate from Sigma, TNFα from R&D systems and cisplatin from Ebewe. To evaluate variability in stress response activation by DILI compounds, PHHs were exposed to a broad concentration range of acetaminophen, propylthiouracyl, nitrofurantoin, ticlopidine, nefazodone and diclofenac (Supporting Table S1) from Sigma. For all compounds except TNF and cisplatin, stocks were prepared in DMSO (BioSolve) at a 500x dilution resulting in a maximum of 0.2% DMSO concentration during exposure. As controls, medium and 0.2% DMSO were taken along.

##### *Cell viability*

To evaluate cytotoxicity, LDH release was evaluated after 24 h of exposure in supernatants using cytotoxicity detection kit from Roche according to instructions by provider. As positive control, supernatant of PHHs incubated for 5 minutes with 1% triton was taken along. Collected supernatants were stored at 4°C for a maximum of 3 days before analysis. Upon analysis, supernatants were diluted 10x and measured in triplicate. Absorbance was measured at 490 nm with a VICTOR plate reader (PerkinElmer). Cell viability was determined for three biological replicates for each condition for each PHH.

##### *Targeted sequencing*

After exposure for 8 or 24 h, the transcriptome was analyzed using the targeted TempO-seq technology (BioSpyder Technologies, Inc., Carlsbad, CA, USA). First, cells were washed with 1x PBS and lysed with 50 µL 1x BNN TempO-seq lysis buffer per well (BioSpyder). Lysates were incubated for 15 min at RT and stored at -80°C. Samples were shipped for TempO-seq analysis (1) using the S1500+ gene set of NIEHS (2) supplemented with additional stress response relevant genes (Supporting Data S2) at BioSpyder (Carlsbad, CA, USA) and sequenced using a HiSeq 2500 Ultra-High-Throughput Sequencing System (Illumina, San Diego, CA, USA). For each PHH, three biological replicates were analyzed. To evaluate variance in dedifferentiation, samples from PHHs in suspension directly upon thawing, snap-frozen liver tissue or PHHs grown in 2D for 24 h derived from 8 individuals were analyzed using the targeted whole transcriptome panel in combination with TempO-seq technology (BioSpyder) (Supporting Data S2).

*Transcriptomics dose-response analysis*

For the analysis of TempO-seq transcriptome data, as a first step reads were aligned by BioSpyder technologies using the TempO-seqR package. Derived raw counts were normalized using the DESeq2 R package (Love et al. 2014) and log2 transformed. A library size cut-off was used of 100.000 counts to eliminate samples having low amount of total counts (Supporting Fig. S2). To evaluate variability in sensitivity for stress response activation, benchmark concentration (BMC) modelling was done using the BMDExpress 2 software developed by Sciome LCC and NIEHS/NTP/EPA (3,4). Dose response modeling was done for each gene and sample using various models (exponential, linear, polynomial, hill and power model). The BMC was defined as the concentration at which 1 standard deviation of increase in gene expression was seen. For each reference compound, the top 50 activated genes across the PHH panel were defined based on both the BMC and the maximal fold change across concentration range (maxFC) (Supporting Data S2). The median BMC and maxFC were calculated based on these top 50 genes for each PHH. To classify PHHs for their sensitivity, a sensitivity score was calculated based on the sum of the ranking of the median BMCs and maxFC at both time points for the top 50 genes for each reference compound. Principal component analysis (PCA) was done using prcomp from the Stats Rpackage. Hierarchical clustering based on Euclidean distance and Wards method was done using Rpackage pheatmap. Data analysis was done using R 3.4.1 and Rstudio with the following Rpackages: DESeq2, pheatmap, ggplot2, data.table, dplyr, reshape2, stats. To evaluate difference in pathway activation between different PHHs, gene set enrichment analysis (GSEA) in combination with Gene Ontology (GO) gene sets v7.1 (5) retrieved from the MSigDB was performed using the maxFC as input in the GSEA software (derived from joint project of UC San Diego and Broad Institute) (6,7). Visualization of GSEA results was done using Cytoscape 3.8.1 software (8) in combination with EnrichmentMap (9) and WordCloud (10).

*Population statistical modelling*

The BMC-maxFC values distributions were modelled in a Bayesian hierarchical framework. We are interested a priori by inter-subject variability; therefore, our primary observational unit was the individual donor. Genes were considered as exchangeable (in the sense that they are independent and identically distributed).

At the level of the $i^{th}$ subject, we assumed that the counts $n_{ik}$ of genes falling in the $k^{th}$ of the $K$ (equal to either two or four) pre-defined clusters followed a multinomial distribution:

$$\boldsymbol{n}_{\mathrm{i}} = (n_{i1}, \ldots, n_{iK}) \sim \mathcal{M}ultinomial(p_{i1}, \ldots, p_{iK}) \quad (1)$$

At the population level, the subjects' multinomial probabilities were softmax-transformed and the corresponding parameters $\boldsymbol{\beta}$ were assumed to be multivariate-normal-distributed around a population mean $\boldsymbol{\mu}$ with covariance matrix $\boldsymbol{\Omega}$.

$$p_{ik} = \frac{\exp(\beta_{ik})}{\sum_{j=1}^{K} \exp(\beta_{ij})} \tag{2}$$

$$\beta_{i1} = 0 \tag{3}$$

$$(\beta_{i2}, \dots, \beta_{iK}) \sim \mathcal{N}_{K-1}(\boldsymbol{\mu}, \boldsymbol{\Omega}) \tag{4}$$

The prior on each element of $\boldsymbol{\mu}$ was a vague normal distribution:

$$\mu_k \sim \mathcal{N}(0, 5) \tag{5}$$

The prior on $\boldsymbol{\Omega}$ was a Cauchy-LKJ (Lewandowski-Kurowicka-Joe) distribution with a Cauchy-distributed diagonal vector of standard deviations $\boldsymbol{\theta}$:

$$\theta_k \sim Cauchy(0, 2.5) \tag{6}$$

and a LJK-distributed prior on the correlation matrix $\boldsymbol{L}$:

$$\mathbf{L} \sim LKJ(3) \tag{7}$$

The *Stan* statistical software was used to obtain a posterior sample of $\boldsymbol{\mu}$, $\boldsymbol{\theta}$, and $\boldsymbol{L}$ values by Hamiltonian Monte Carlo simulations.

Still at the $i^{th}$ subject level, we modelled independently for each cluster the joint distribution of genes' BMCs (noted x in the following) and maxFCs (noted y) values as a bivariate lognormal distribution. We took the absolute value of negative maxFC values before log-transformation:

$$(\log(\boldsymbol{x}_i), \log(abs(\boldsymbol{y}_i))) \sim \mathcal{N}_2\left((\upsilon_{xi}, \upsilon_{yi}), \boldsymbol{\Delta}_i\right) \tag{8}$$

with $\boldsymbol{\Delta}$ defined as

$$\boldsymbol{\Delta}_i = \begin{pmatrix} \sigma_{xi} & \rho_i \\ \rho_i & \sigma_{yi} \end{pmatrix} \tag{9}$$

At the population level, we modeled the distributions of subjects' means $\upsilon_{xi}$ and $\upsilon_{yi}$, of the standard deviations $\sigma_{xi}$ and $\sigma_{yi}$, and of the correlation coefficient $\rho_i$ as normal around their population counterparts:

$$\upsilon_{xi} \sim \mathcal{N}(\upsilon_x, \sigma_a) \tag{10}$$

$$\upsilon_{yi} \sim \mathcal{N}(\upsilon_y, \sigma_b) \tag{11}$$

$$\sigma_{xi} \sim \mathcal{N}(\sigma_x, \sigma_c) \tag{12}$$

$$\sigma_{yi} \sim \mathcal{N}(\sigma_y, \sigma_d) \tag{13}$$

$$\rho_i \sim \mathcal{N}(\rho, \sigma_e) \tag{14}$$

The standard deviations $\sigma_a$, $\sigma_b$, $\sigma_c$, $\sigma_d$, $\sigma_e$, were all assigned a half normal prior with SD 0.2. The other priors were:

$$\upsilon_x \sim \mathcal{U}niform(-5,7) \tag{15}$$

$$\upsilon_y \sim \mathcal{U}niform\,(-11.5, 5) \tag{16}$$

$$\sigma_x \sim \mathcal{H}alfnormal(1) \tag{17}$$

$$\sigma_y \sim \mathcal{H}alfnormal(1) \tag{18}$$

$$\rho \sim \mathcal{U}niform\,(-1,1) \tag{19}$$

All model parameters (265 parameters in total for each chemical exposure) were jointly estimated from the data with Metropolis-Hastings Markov chain Monte Carlo simulation, using the *GNU MCSim* software V6.1.0.

*Predictive simulations*

For predictions of inter-subject variability, the transformed BMC and maxFC values were restored to natural space using the suitable inverse transformations. First, large sample reference median coefficients of variation (CVs) were obtained for BMC and maxFC values by simulating of 1000 assays with hepatocytes from 2000 individuals each. For each simulated individual, BMC and maxFC values were simulated by Monte Carlo sampling using the average posterior estimates of the population parameters obtained by calibration of the model with the experimental data on 50 individuals. Those can therefore be considered as "true CV values", conditionally on the BMC-maxFC values being correctly modelled. Similar assay simulations were performed for smaller, realistic, donor panel sizes (N = 3, 4, 5, 6, 7, 8, 10, 12, 14, 16, 20, 24, 28, 32, 36, 40, 50). CVs for BMC and maxFC values were obtained for each case.

#### Supporting Figure Legends

**Supporting Fig. S1. Characteristics of large panel of primary human hepatocytes derived from 50 individuals.** Distribution of gender (A), age (B) and BMI (C) within panel of primary human hepatocytes derived from 50 individuals.

**Supporting Fig. S2. Quality controls of TempO-seq data for each PHH donor.** A) Distribution of total library size for each sample across PHH panel. B) PearsonR correlation of each biological replicate compared to mean for each sample across PHH panel. N=3

**Supporting Fig. S3. Principal component analysis (PCA) of stress responsive genes at basal conditions across panel of PHHs.** A) PCA based on log2 normalized counts of stress responsive genes of panel of PHHs at basal medium conditions at identical timepoints of compound exposure for 8 and 24 h. B) Top 5 genes mostly determining PC1 or 2 depicted as vectors representing contribution and PC orientation. N=3.

**Supporting Fig. S4. Variability in expression of liver-related genes across PHH panel.** A) Hierarchical clustering of log2 normalized counts of liver-related genes within the S1500+ gene set across PHH panel derived from 50 individuals cultured as 2D for 24 h upon thawing. B) Hierarchical clustering of log2 normalized counts of liver-related genes within the targeted whole transcriptome gene set for liver tissue (N=1), freshly thawed PHHs in suspension (N=3) or PHHs cultured as 2D for 24 h (N=3) derived from 8 individuals.

**Supporting Fig. S5. Inter-individual variability of median maximal fold change and bench mark concentration of most responsive genes upon chemical exposure.** A) Hierarchical clustering of relative median maximal fold change (maxFC) across concentration range for top 50 stress responsive genes for each compound across PHH panel at 8 and 24 h exposure. B) Hierarchical clustering of relative median BMC of top 50 stress responsive genes for each compound and timepoint across PHH panel. N=3.

**Supporting Fig. S6. Variability in the distribution of maximal fold change of most responsive genes upon exposure.** Distribution of maximal fold change across concentration range (maxFC) of top 50 stress responsive genes for each compound (diethyl maleate; DEM, cisplatin; CPT, tunicamycin; TUN, TNFα; TNF) and time point 8 and 24 h. Lines represent each PHH where colour reflects the sensitivity score based on rank of median maxFC and bench mark concentration (BMC) of top 50 genes.

**Supporting Fig. S7. Correlation cell viability and sensitivity scores for PHH panel.** Correlation between the sensitivity scores for each compound and LDH leakage as a measure of cell viability at 3300 µM diethyl maleate (DEM). The sensitivity scores were based on rank of median maxFC and bench mark concentration (BMC) of top 50 stress responsive genes for each compound (diethyl maleate; DEM, cisplatin; CPT, tunicamycin; TUN, TNFα; TNF) ranging from 0 (most sensitive) to 200 (most insensitive). Left bottom represents the scatter plots, in the middle histograms and upper right the Pearson correlation. Significant correlations depicted as * $p < 0.05$, ** $p < 0.01$, *** $p < 0.001$. N = 3.

**Supporting Fig. S8. Effect of pathology background on sensitivity of chemical-induced stress response activation.** Distribution of median benchmark concentration (BMC) (A) or median maximal fold change across concentration range (maxFC) (B) of top 50 stress responsive genes for each compound (diethyl maleate; DEM, cisplatin; CPT, tunicamycin; TUN, TNFα; TNF) and time point 8 and 24 h across PHH panel from 50 individuals with or without cancer. C) Distribution of normalized counts in baseline medium condition of top 50 stress responsive genes for each compound and time point across PHH panel from 50 individuals with or without liver pathology (left panel) or cancer (right panel). Significance levels represented as * $p < 0.1$, ** $p < 0.05$, *** $p < 0.01$. N = 3.

**Supporting Fig. S9. Correlation hepatic phenotype with sensitivity of chemical-induced stress response activation.** Scatter plots of the median expression of liver-related genes vs median maximal fold change across concentration range (maxFC) (left panel) or median benchmark concentration (BMC) (right panel) of top 50 stress responsive genes for each compound (diethyl maleate; DEM, cisplatin; CPT, tunicamycin; TUN, TNFα; TNF) and time point 8 and 24 h for each PHH across panel of 50. Sensitivity scores for each PHH is depicted in color scale from red to blue, sensitive to insensitive. The correlation is given as $r^2$.

**Supporting Fig. S10. Evaluation of variance in pathway activation by gene set enrichment analysis (GSEA) of gene ontology (GO) terms.** Gene set enrichment analysis (GSEA) (6) was done using gene ontology (GO) terms v7.1 and the maximal fold change across concentration range (maxFC) of measured S1500+ genes as input for each PHH and treatment for 24h with cisplatin (CPT), diethyl maleate (DEM), TNFα (TNF) or tunicamycin (TUN). Detailed representation of significantly enriched GO terms in which one or more terms within cluster showed specific enrichment in either most sensitive (red circle) or insensitive (blue circle) PHHs, or both defined as core (yellow circle) using Cytoscape (8) and EnrichmentMap (9). Within each term, the NES for each PHH is depicted in grey to red scale from low to high.

Enriched GO terms were clustered and summarized with 3 to 4 keywords using WordCloud (10).

**Supporting Fig. S11. Simulation of the distribution of bench mark concentration (BMC) and maximal fold change across concentration range (maxFC) of top 50 stress responsive genes across extended panel of PHHs.** A) Experimental observation of distribution of BMC and maxFC of top 50 stress responsive genes for diethyl maleate (DEM), cisplatin (CPT), tunicamycin (TUN) and TNFα (TNF) across panel of PHHs depicted in different colours at 8 and 24 h timepoint. N = 3. B) Computer simulated sample for a panel of 500 PHHs depicted in different colours. The vertical and horizontal lines delimit the observed cluster of BMC and maxFC values.

**Supporting Fig. S12. Simulation of the confidence intervals of the lognormal bivariate distributions of bench mark concentration (BMC) and maximal fold change across concentration range (maxFC) of top 50 stress responsive genes across panel of PHHs.** Experimental observation of the distribution of BMC and maxFC of top 50 stress responsive genes for diethyl maleate (DEM), cisplatin (CPT), tunicamycin (TUN) and TNFα (TNF) across panel of PHHs depicted as dots in different colours at 8 (upper panel) and 24 h (lower panel) timepoint for each observed section or quadrant. N = 3. Computer simulated confidence intervals indicated as eclipses for a panel of 500 PHHs depicted in different colours.

#### Supporting Tables

**Supporting Table S1. Overview of concentration ranges used for each compound tested in panel of PHHs.**

#### Supporting Data

**Supporting Data S1. Information PHH panel from 50 individuals.**

**Supporting Data S2. List of S1500+ genes, whole transcriptome and top 50 most responsive genes**

**Supporting Data S3. BMC and maxFC for top 50 most responsive genes**

**Supporting Data S4. Sensitivity scores for PHH panel**

**Supporting Data S5. Gene set enrichment analysis (GSEA) of gene ontology (GO) terms based on maximal fold change across concentration range for panel of primary human hepatocytes.** Enriched pathways were specified being enriched in only sensitive or insensitive PHHs or enriched in both defined as core pathways. Top 10 enriched pathways are depicted based on mean FDR.

**Supporting Table S1. Overview of concentration ranges used for each compound tested in panel of PHHs.**

| *ID* | ***Reference compounds*** | | | |
|---|---|---|---|---|
| | **DEM** | **CPT** | **TUN** | **TNF** |
| *1* | 1 | 0.1 | 0.0001 | 0.1 |
| *2* | 10 | 1 | 0.001 | 0.3 |
| *3* | 100 | 3.3 | 0.01 | 1 |
| *4* | 330 | 10 | 0.1 | 3.3 |
| *5* | 1000 | 33 | 1 | 10 |
| *6* | 3300 | 100 | 10 | 33 |
| | µM | µM | µM | ng/mL |

| *xCmax* | ***DILI compounds*** | | | | | |
|---|---|---|---|---|---|---|
| | **NIT** | **ACE** | **PRO** | **TIC** | **DIC** | **NEF** |
| *1* | 6 | 139 | 9.1 | 8.1 | 10.1 | 4.0 |
| *2.5* | 15 | 347.5 | 22.75 | 20.2 | 25.25 | 9.9 |
| *5* | 30 | 695 | 45.5 | 40.4 | 50.5 | 19.8 |
| *10* | 60 | 1390 | 91 | 80.7 | 101 | 39.5 |
| *25* | 150 | 3475 | 227.5 | 201.9 | 252.5 | 98.8 |
| *50* | 300 | 6950 | 455 | 403.7 | 505 | 197.5 |
| *100* | 600 | 13900 | 910 | 807.5 | 1010 | 395.0 |
| | µM | µM | µM | µM | µM | µM |

*Diethyl maleate; DEM, Cisplatin; CPT, tunicamycin; TUN, TNFα; TNF, acetaminophen; ACE, propylthiouracyl; PRO, nitrofurantoin; NIT, ticlopidine; TIC, nefazodone; NEF, diclofenac; DIC.*

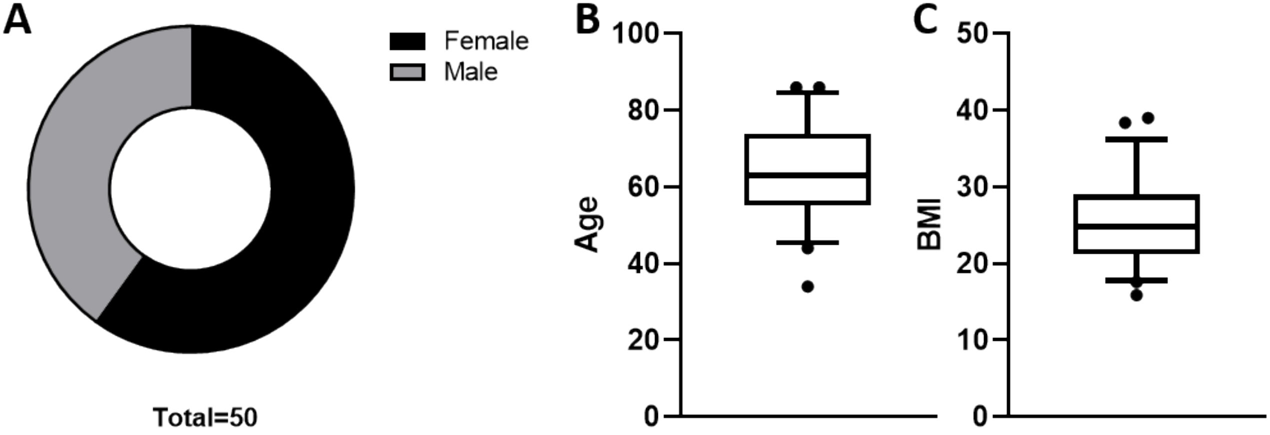
A
Female
Male
Total=50
B
100
80
60
40
20
0
Age
C
50
40
30
20
10
0
BMI

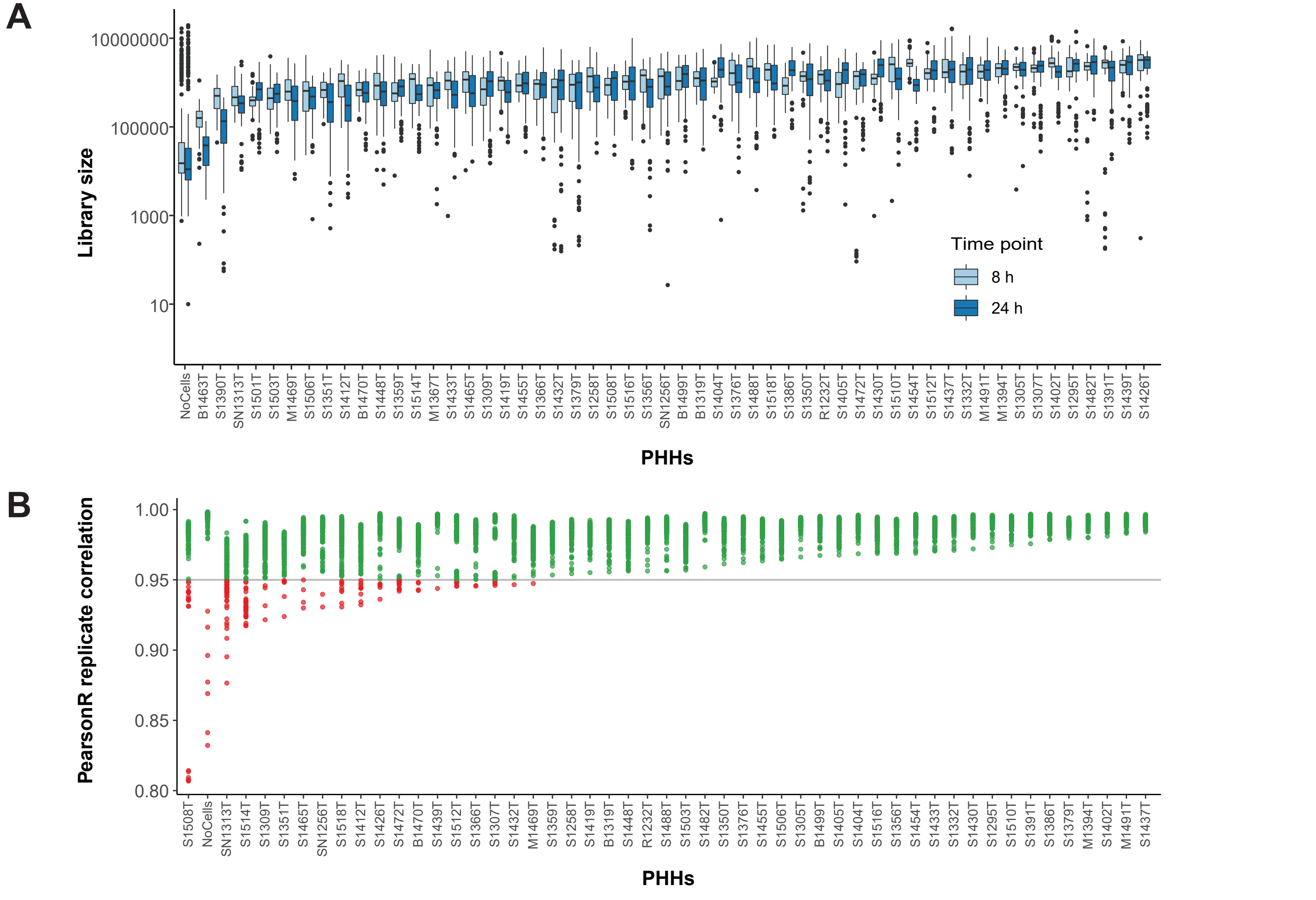

A
Library size
10000000
100000
1000
10
Time point
8 h
24 h
NoCells B1463T S1390T SN1313T S1501T S1503T M1469T S1506T S1351T S1412T B1470T S1448T S1359T S1514T M1367T S1433T S1465T S1309T S1419T S1455T S1366T S1432T S1379T S1258T S1508T S1516T S1356T SN1256T B1499T B1319T S1404T S1376T S1488T S1518T S1386T S1350T R1232T S1405T S1472T S1430T S1510T S1454T S1512T S1437T S1332T M1491T M1394T S1305T S1307T S1402T S1295T S1482T S1391T S1439T S1426T
PHHs
B
PearsonR replicate correlation
1.00
0.95
0.90
0.85
0.80
S1508T NoCells SN1313T S1514T S1309T S1351T S1465T SN1256T S1518T S1412T S1426T S1472T B1470T S1439T S1512T S1366T S1307T S1432T M1469T S1359T S1258T S1419T B1319T S1448T R1232T S1488T S1503T S1482T S1350T S1376T S1455T S1506T S1305T B1499T S1405T S1404T S1516T S1356T S1454T S1433T S1332T S1430T S1295T S1510T S1391T S1386T S1379T M1394T S1402T M1491T S1437T
PHHs

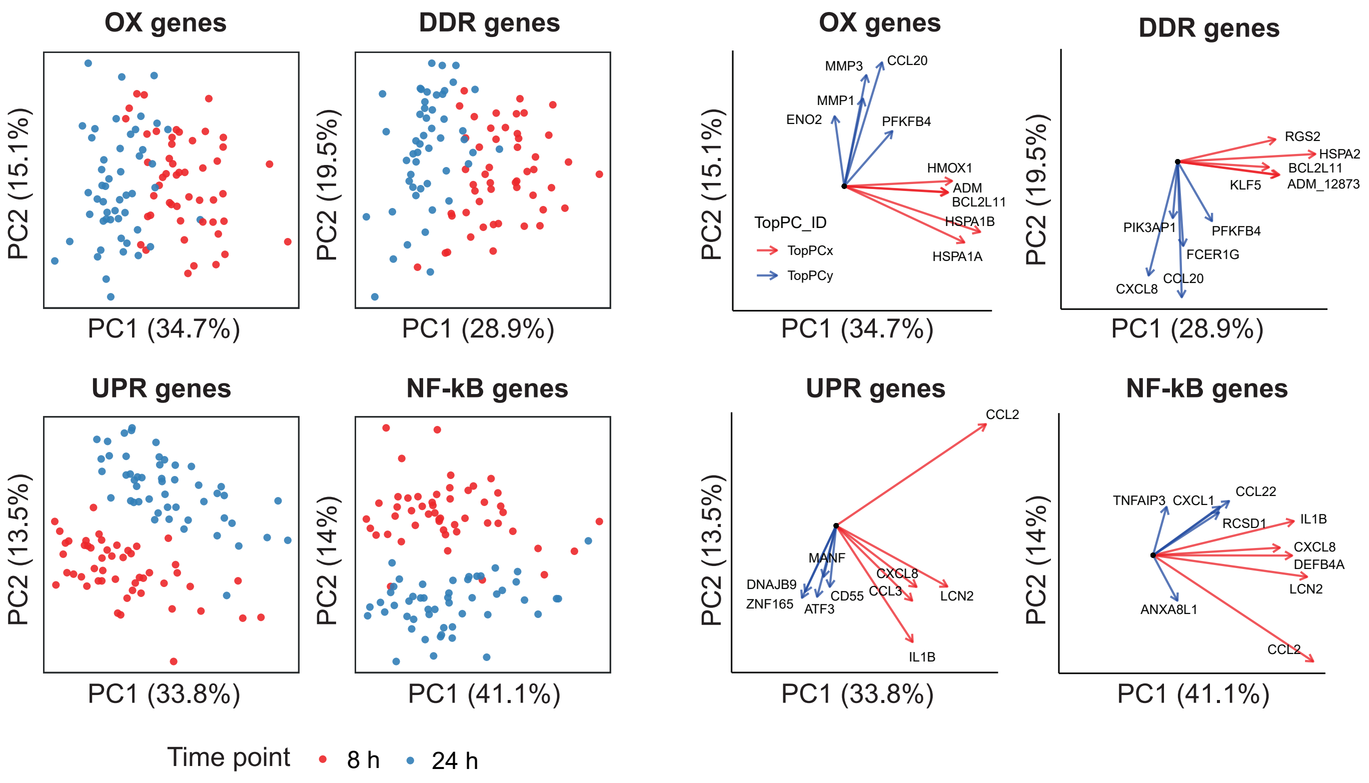

OX genes
DDR genes
PC2 (15.1%)
PC1 (34.7%)
PC2 (19.5%)
PC1 (28.9%)
UPR genes
NF-kB genes
PC2 (13.5%)
PC1 (33.8%)
PC2 (14%)
PC1 (41.1%)
Time point
8 h
24 h
OX genes
MMP3
CCL20
MMP1
ENO2
PFKFB4
HMOX1
ADM
BCL2L11
HSPA1B
HSPA1A
TopPC_ID
TopPCx
TopPCy
DDR genes
RGS2
HSPA2
BCL2L11
KLF5
ADM_12873
PIK3AP1
PFKFB4
FCER1G
CCL20
CXCL8
UPR genes
CCL2
MANF
DNAJB9
ZNF165
ATF3
CD55
CXCL8
CCL3
LCN2
IL1B
NF-kB genes
CCL22
TNFAIP3
CXCL1
RCSD1
IL1B
CXCL8
DEFB4A
LCN2
ANXA8L1
CCL2

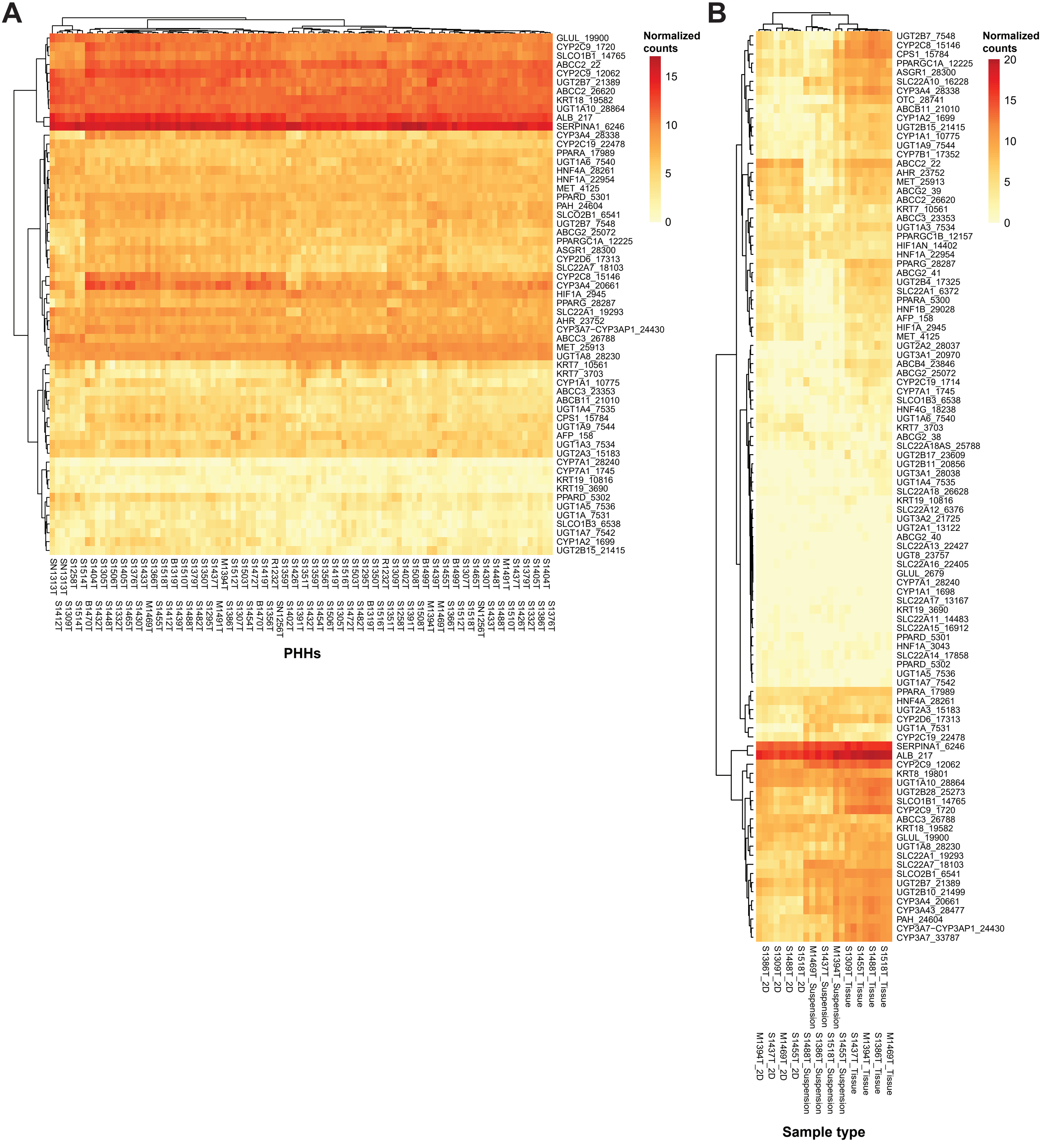
A
GLUL_19900
CYP2C9_1720
SLCO1B1_14765
ABCC2_22
CYP2C9_12062
UGT2B7_21389
ABCC2_26620
KRT18_19582
UGT1A10_28864
ALB_217
SERPINA1_6246
CYP3A4_28338
CYP2C19_22478
PPARA_17989
UGT1A6_7540
HNF4A_28261
HNF1A_22954
MET_4125
PPARD_5301
PAH_24604
SLCO2B1_6541
UGT2B7_7548
ABCG2_25072
PPARGC1A_12225
ASGR1_28300
CYP2D6_17313
SLC22A7_18103
CYP2C8_15146
CYP3A4_20661
HIF1A_2945
PPARG_28287
SLC22A1_19293
AHR_23752
CYP3A7−CYP3AP1_24430
ABCC3_26788
MET_25913
UGT1A8_28230
KRT7_10561
KRT7_3703
CYP1A1_10775
ABCC3_23353
ABCB11_21010
UGT1A4_7535
CPS1_15784
UGT1A9_7544
AFP_158
UGT1A3_7534
UGT2A3_15183
CYP7A1_28240
CYP7A1_1745
KRT19_10816
KRT19_3690
PPARD_5302
UGT1A5_7536
UGT1A_7531
SLCO1B3_6538
UGT1A7_7542
CYP1A2_1699
UGT2B15_21415
Normalized counts
15
10
5
0
PHHs
B
UGT2B7_7548
CYP2C8_15146
CPS1_15784
PPARGC1A_12225
ASGR1_28300
SLC22A10_16228
CYP3A4_28338
OTC_28741
ABCB11_21010
CYP1A2_1699
UGT2B15_21415
CYP1A1_10775
UGT1A9_7544
CYP7B1_17352
ABCC2_22
AHR_23752
MET_25913
ABCG2_39
ABCC2_26620
KRT7_10561
ABCC3_23353
UGT1A3_7534
PPARGC1B_12157
HIF1AN_14402
HNF1A_22954
PPARG_28287
ABCG2_41
UGT2B4_17325
SLC22A1_6372
PPARA_5300
HNF1B_29028
AFP_158
HIF1A_2945
MET_4125
UGT2A2_28037
UGT3A1_20970
ABCB4_23846
ABCG2_25072
CYP2C19_1714
CYP7A1_1745
SLCO1B3_6538
HNF4G_18238
UGT1A6_7540
KRT7_3703
ABCG2_38
SLC22A18AS_25788
UGT2B17_23609
UGT2B11_20856
UGT3A1_28038
UGT1A4_7535
SLC22A18_26628
KRT19_10816
SLC22A12_6376
UGT3A2_21725
UGT2A1_13122
ABCG2_40
SLC22A13_22427
UGT8_23757
SLC22A16_22405
GLUL_2679
CYP7A1_28240
CYP1A1_1698
SLC22A17_13167
KRT19_3690
SLC22A11_14483
SLC22A15_16912
PPARD_5301
HNF1A_3043
SLC22A14_17858
PPARD_5302
UGT1A5_7536
UGT1A7_7542
PPARA_17989
HNF4A_28261
UGT2A3_15183
CYP2D6_17313
UGT1A_7531
CYP2C19_22478
SERPINA1_6246
ALB_217
CYP2C9_12062
KRT8_19801
UGT1A10_28864
UGT2B28_25273
SLCO1B1_14765
CYP2C9_1720
ABCC3_26788
KRT18_19582
GLUL_19900
UGT1A8_28230
SLC22A1_19293
SLC22A7_18103
SLCO2B1_6541
UGT2B7_21389
UGT2B10_21499
CYP3A4_20661
CYP3A43_28477
PAH_24604
CYP3A7−CYP3AP1_24430
CYP3A7_33787
Normalized counts
20
15
10
5
0
M1394T_2D
S1386T_2D
S1437T_2D
S1309T_2D
M1469T_2D
S1488T_2D
S1455T_2D
S1518T_2D
S1488T_Suspension
M1469T_Suspension
S1386T_Suspension
S1437T_Suspension
S1518T_Suspension
M1394T_Suspension
S1455T_Suspension
S1309T_Tissue
S1437T_Tissue
S1455T_Tissue
M1394T_Tissue
S1488T_Tissue
S1386T_Tissue
S1518T_Tissue
M1469T_Tissue
Sample type

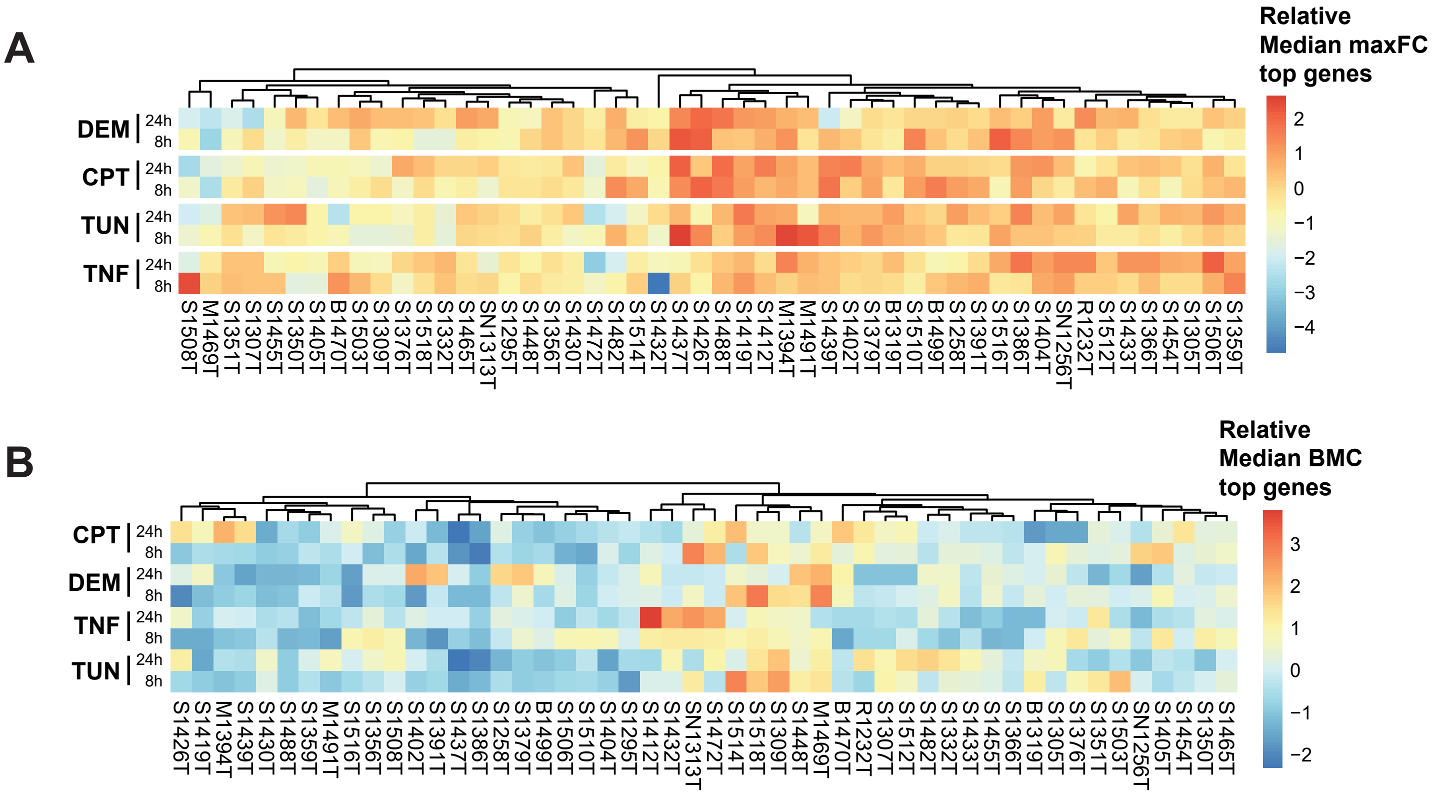

A
Relative
Median maxFC
top genes
DEM
CPT
TUN
TNF
24h
8h
2
1
0
−1
−2
−3
−4
S1508T
M1469T
S1351T
S1307T
S1455T
S1350T
S1405T
B1470T
S1503T
S1309T
S1376T
S1518T
S1332T
S1465T
SN1313T
S1295T
S1448T
S1356T
S1430T
S1472T
S1482T
S1514T
S1432T
S1437T
S1426T
S1488T
S1419T
S1412T
M1394T
M1491T
S1439T
S1402T
S1379T
B1319T
S1510T
B1499T
S1258T
S1391T
S1516T
S1386T
S1404T
SN1256T
R1232T
S1512T
S1433T
S1366T
S1454T
S1305T
S1506T
S1359T
B
Relative
Median BMC
top genes
CPT
DEM
TNF
TUN
24h
8h
3
2
1
0
−1
−2
S1426T
S1419T
M1394T
S1439T
S1430T
S1488T
S1359T
M1491T
S1516T
S1356T
S1508T
S1402T
S1391T
S1437T
S1386T
S1258T
S1379T
B1499T
S1506T
S1510T
S1404T
S1295T
S1412T
S1432T
SN1313T
S1472T
S1514T
S1518T
S1309T
S1448T
M1469T
B1470T
R1232T
S1307T
S1512T
S1482T
S1332T
S1433T
S1455T
S1366T
B1319T
S1305T
S1376T
S1351T
S1503T
SN1256T
S1405T
S1454T
S1350T
S1465T

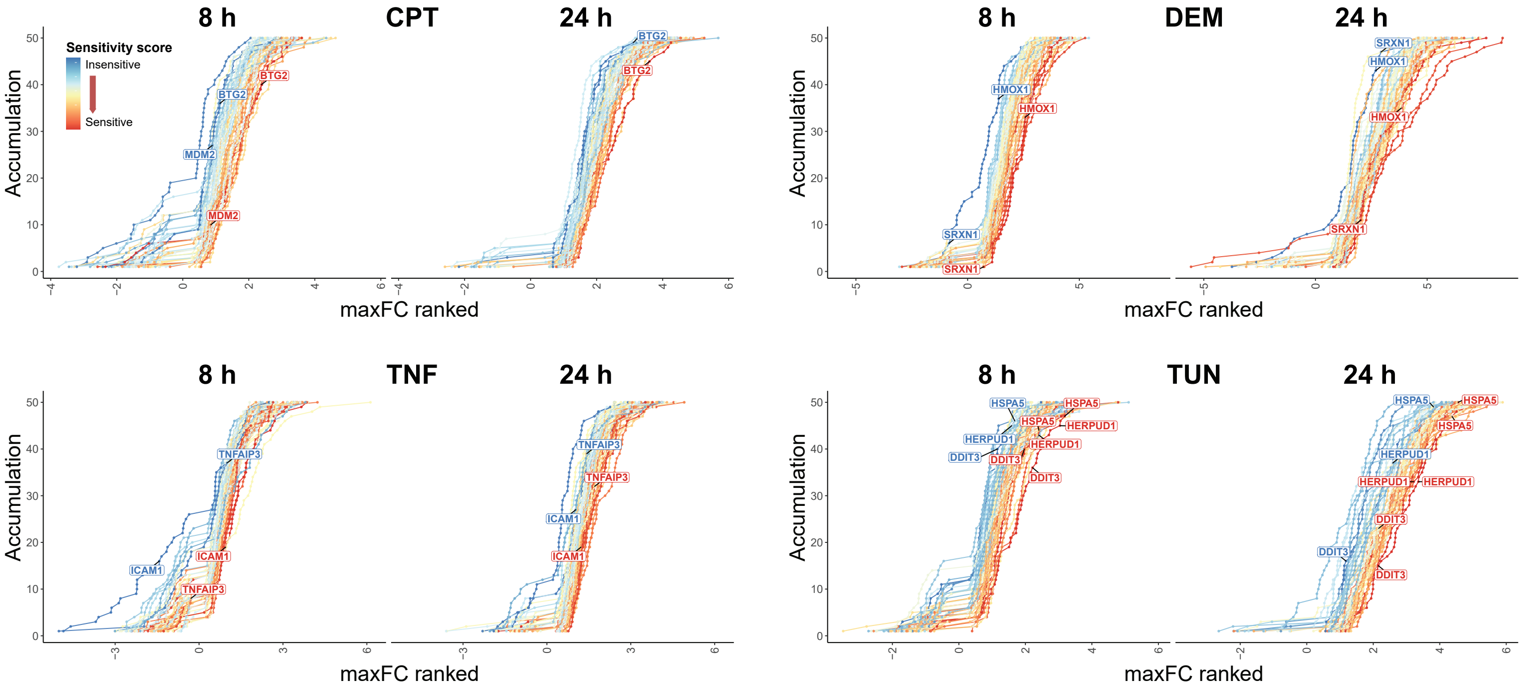
CPT
8 h
24 h
Sensitivity score
Insensitive
Sensitive
BTG2
MDM2
Accumulation
maxFC ranked
DEM
HMOX1
SRXN1
TNF
TNFAIP3
ICAM1
TUN
HSPA5
HERPUD1
DDIT3

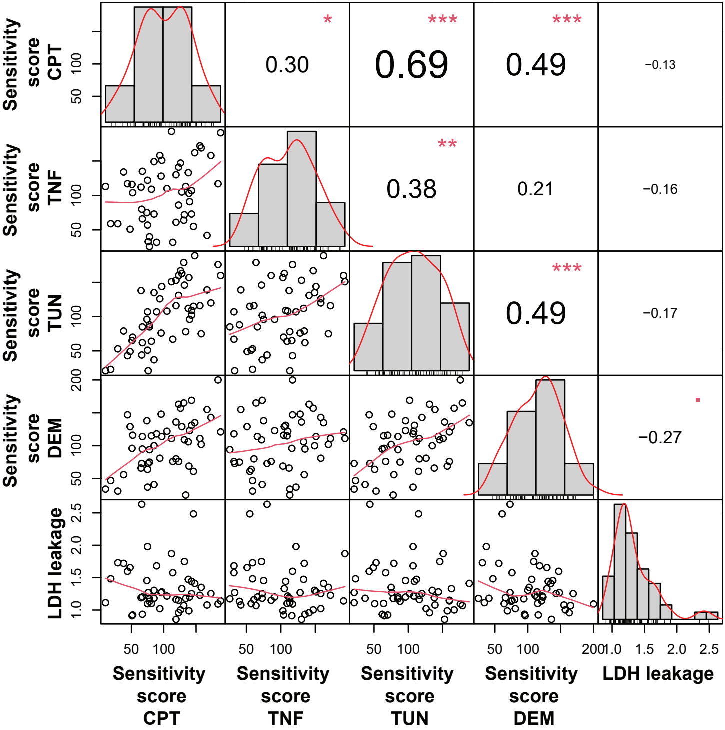

Sensitivity score CPT
Sensitivity score TNF
Sensitivity score TUN
Sensitivity score DEM
LDH leakage
0.30
0.69
0.49
−0.13
0.38
0.21
−0.16
0.49
−0.17
−0.27
*
***
***
**
***

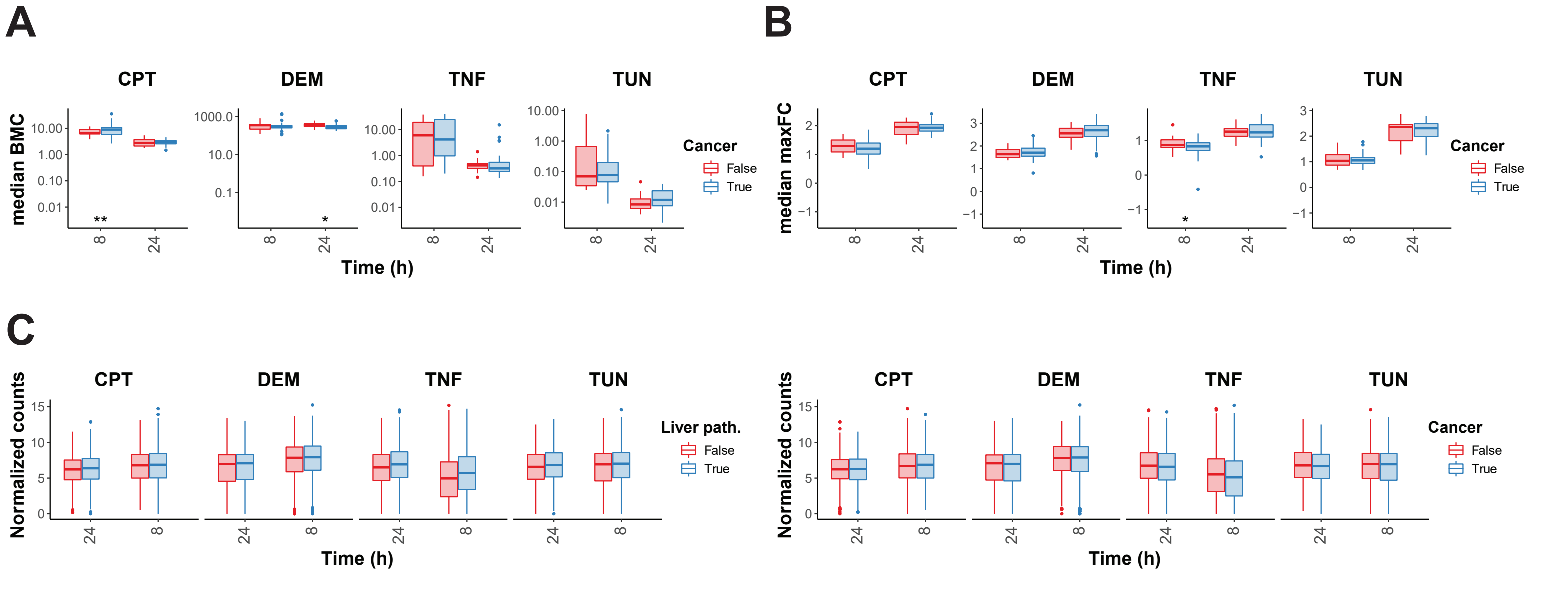
A
B
C
CPT
DEM
TNF
TUN
median BMC
median maxFC
Normalized counts
Time (h)
Cancer
Liver path.
False
True

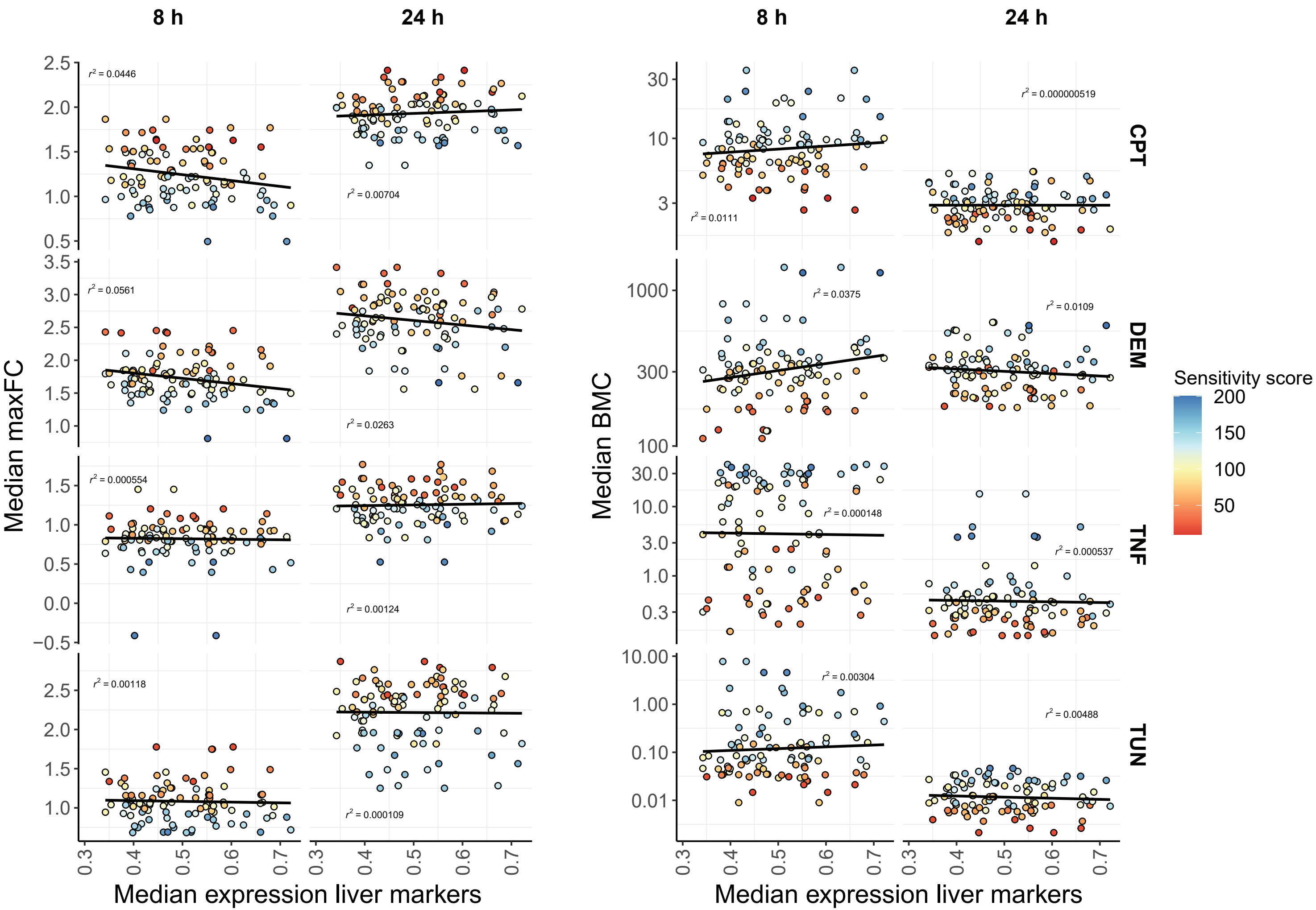

8 h
24 h
8 h
24 h
Median maxFC
Median BMC
Median expression liver markers
Median expression liver markers
CPT
DEM
TNF
TUN
Sensitivity score
200
150
100
50
r² = 0.0446
r² = 0.00704
r² = 0.0111
r² = 0.000000519
r² = 0.0561
r² = 0.0263
r² = 0.0375
r² = 0.0109
r² = 0.000554
r² = 0.00124
r² = 0.000148
r² = 0.000537
r² = 0.00118
r² = 0.000109
r² = 0.00304
r² = 0.00488

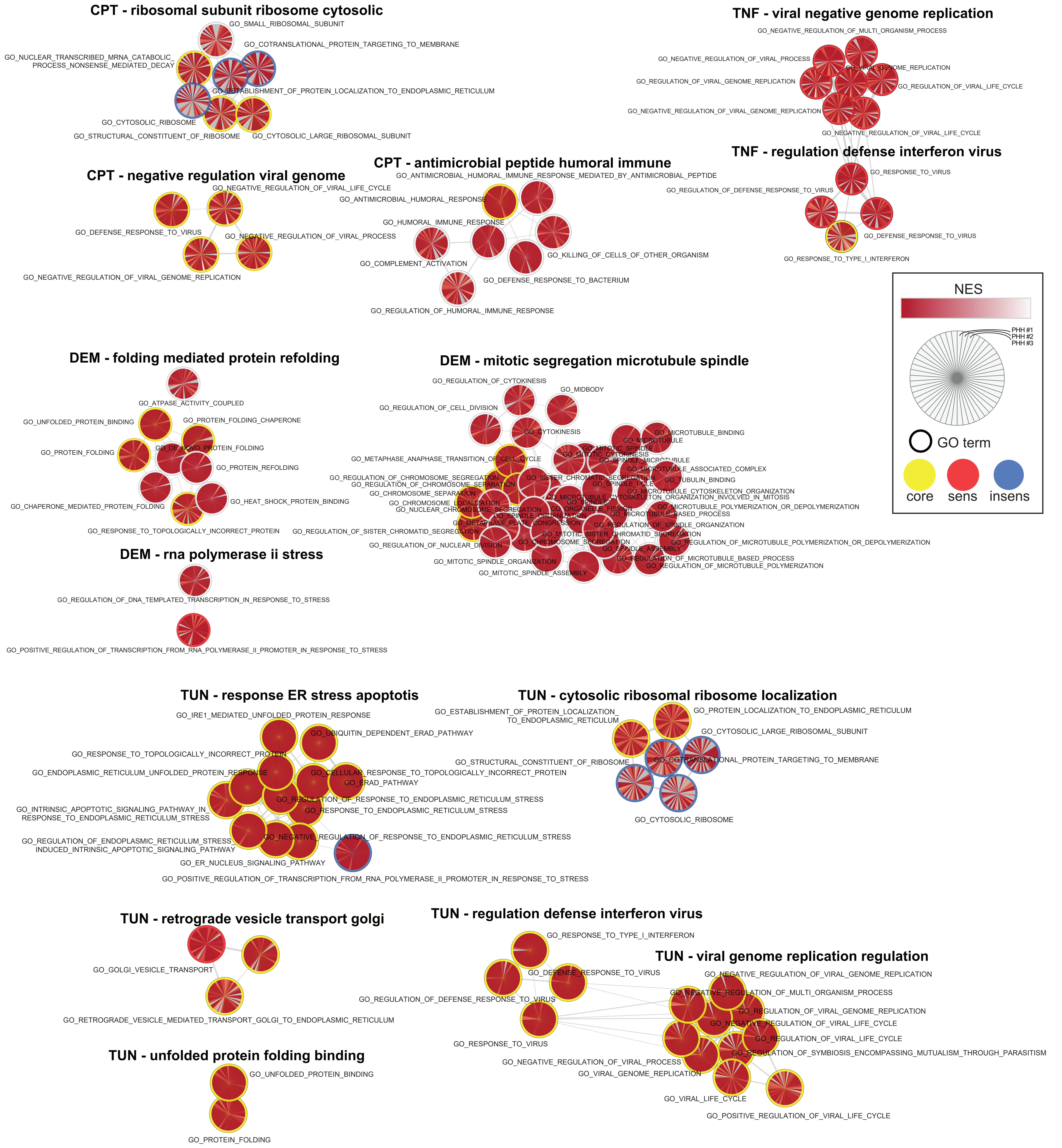

CPT - ribosomal subunit ribosome cytosolic
GO_SMALL_RIBOSOMAL_SUBUNIT
GO_COTRANSLATIONAL_PROTEIN_TARGETING_TO_MEMBRANE
GO_NUCLEAR_TRANSCRIBED_MRNA_CATABOLIC_
PROCESS_NONSENSE_MEDIATED_DECAY
GO_ESTABLISHMENT_OF_PROTEIN_LOCALIZATION_TO_ENDOPLASMIC_RETICULUM
GO_CYTOSOLIC_RIBOSOME
GO_STRUCTURAL_CONSTITUENT_OF_RIBOSOME
GO_CYTOSOLIC_LARGE_RIBOSOMAL_SUBUNIT
TNF - viral negative genome replication
GO_NEGATIVE_REGULATION_OF_MULTI_ORGANISM_PROCESS
GO_NEGATIVE_REGULATION_OF_VIRAL_PROCESS
GO_VIRAL_GENOME_REPLICATION
GO_REGULATION_OF_VIRAL_GENOME_REPLICATION
GO_REGULATION_OF_VIRAL_LIFE_CYCLE
GO_NEGATIVE_REGULATION_OF_VIRAL_GENOME_REPLICATION
GO_NEGATIVE_REGULATION_OF_VIRAL_LIFE_CYCLE
CPT - antimicrobial peptide humoral immune
GO_ANTIMICROBIAL_HUMORAL_IMMUNE_RESPONSE_MEDIATED_BY_ANTIMICROBIAL_PEPTIDE
GO_ANTIMICROBIAL_HUMORAL_RESPONSE
GO_HUMORAL_IMMUNE_RESPONSE
GO_KILLING_OF_CELLS_OF_OTHER_ORGANISM
GO_COMPLEMENT_ACTIVATION
GO_DEFENSE_RESPONSE_TO_BACTERIUM
GO_REGULATION_OF_HUMORAL_IMMUNE_RESPONSE
CPT - negative regulation viral genome
GO_NEGATIVE_REGULATION_OF_VIRAL_LIFE_CYCLE
GO_DEFENSE_RESPONSE_TO_VIRUS
GO_NEGATIVE_REGULATION_OF_VIRAL_PROCESS
GO_NEGATIVE_REGULATION_OF_VIRAL_GENOME_REPLICATION
TNF - regulation defense interferon virus
GO_RESPONSE_TO_VIRUS
GO_REGULATION_OF_DEFENSE_RESPONSE_TO_VIRUS
GO_DEFENSE_RESPONSE_TO_VIRUS
GO_RESPONSE_TO_TYPE_I_INTERFERON
NES
PHH #1
PHH #2
PHH #3
GO term
core
sens
insens
DEM - folding mediated protein refolding
GO_ATPASE_ACTIVITY_COUPLED
GO_UNFOLDED_PROTEIN_BINDING
GO_PROTEIN_FOLDING_CHAPERONE
GO_DE_NOVO_PROTEIN_FOLDING
GO_PROTEIN_FOLDING
GO_PROTEIN_REFOLDING
GO_CHAPERONE_MEDIATED_PROTEIN_FOLDING
GO_HEAT_SHOCK_PROTEIN_BINDING
GO_RESPONSE_TO_TOPOLOGICALLY_INCORRECT_PROTEIN
DEM - mitotic segregation microtubule spindle
GO_REGULATION_OF_CYTOKINESIS
GO_MIDBODY
GO_REGULATION_OF_CELL_DIVISION
GO_CYTOKINESIS
GO_MICROTUBULE_BINDING
GO_MICROTUBULE
GO_MITOTIC_SPINDLE
GO_MITOTIC_CYTOKINESIS
GO_METAPHASE_ANAPHASE_TRANSITION_OF_CELL_CYCLE
GO_SPINDLE_MICROTUBULE
GO_MICROTUBULE_ASSOCIATED_COMPLEX
GO_REGULATION_OF_CHROMOSOME_SEGREGATION
GO_SISTER_CHROMATID_SEGREGATION
GO_TUBULIN_BINDING
GO_REGULATION_OF_CHROMOSOME_SEPARATION
GO_SPINDLE_POLE
GO_CHROMOSOME_SEPARATION
GO_MICROTUBULE_CYTOSKELETON_ORGANIZATION
GO_MICROTUBULE_CYTOSKELETON_ORGANIZATION_INVOLVED_IN_MITOSIS
GO_CHROMOSOME_LOCALIZATION
GO_SPINDLE
GO_MICROTUBULE_POLYMERIZATION_OR_DEPOLYMERIZATION
GO_NUCLEAR_CHROMOSOME_SEGREGATION
GO_ORGANELLE_FISSION
GO_SPINDLE_ORGANIZATION
GO_MICROTUBULE_BASED_PROCESS
GO_METAPHASE_PLATE_CONGRESSION
GO_REGULATION_OF_SPINDLE_ORGANIZATION
GO_REGULATION_OF_SISTER_CHROMATID_SEGREGATION
GO_MITOTIC_SISTER_CHROMATID_SEGREGATION
GO_CHROMOSOME_SEGREGATION
GO_REGULATION_OF_MICROTUBULE_POLYMERIZATION_OR_DEPOLYMERIZATION
GO_REGULATION_OF_NUCLEAR_DIVISION
GO_SPINDLE_ASSEMBLY
GO_REGULATION_OF_MICROTUBULE_BASED_PROCESS
GO_MITOTIC_SPINDLE_ORGANIZATION
GO_REGULATION_OF_MICROTUBULE_POLYMERIZATION
GO_MITOTIC_SPINDLE_ASSEMBLY
DEM - rna polymerase ii stress
GO_REGULATION_OF_DNA_TEMPLATED_TRANSCRIPTION_IN_RESPONSE_TO_STRESS
GO_POSITIVE_REGULATION_OF_TRANSCRIPTION_FROM_RNA_POLYMERASE_II_PROMOTER_IN_RESPONSE_TO_STRESS
TUN - response ER stress apoptotis
GO_IRE1_MEDIATED_UNFOLDED_PROTEIN_RESPONSE
GO_UBIQUITIN_DEPENDENT_ERAD_PATHWAY
GO_RESPONSE_TO_TOPOLOGICALLY_INCORRECT_PROTEIN
GO_ENDOPLASMIC_RETICULUM_UNFOLDED_PROTEIN_RESPONSE
GO_CELLULAR_RESPONSE_TO_TOPOLOGICALLY_INCORRECT_PROTEIN
GO_ERAD_PATHWAY
GO_INTRINSIC_APOPTOTIC_SIGNALING_PATHWAY_IN_
RESPONSE_TO_ENDOPLASMIC_RETICULUM_STRESS
GO_REGULATION_OF_RESPONSE_TO_ENDOPLASMIC_RETICULUM_STRESS
GO_RESPONSE_TO_ENDOPLASMIC_RETICULUM_STRESS
GO_REGULATION_OF_ENDOPLASMIC_RETICULUM_STRESS_
INDUCED_INTRINSIC_APOPTOTIC_SIGNALING_PATHWAY
GO_NEGATIVE_REGULATION_OF_RESPONSE_TO_ENDOPLASMIC_RETICULUM_STRESS
GO_ER_NUCLEUS_SIGNALING_PATHWAY
GO_POSITIVE_REGULATION_OF_TRANSCRIPTION_FROM_RNA_POLYMERASE_II_PROMOTER_IN_RESPONSE_TO_STRESS
TUN - cytosolic ribosomal ribosome localization
GO_ESTABLISHMENT_OF_PROTEIN_LOCALIZATION_
TO_ENDOPLASMIC_RETICULUM
GO_PROTEIN_LOCALIZATION_TO_ENDOPLASMIC_RETICULUM
GO_CYTOSOLIC_LARGE_RIBOSOMAL_SUBUNIT
GO_STRUCTURAL_CONSTITUENT_OF_RIBOSOME
GO_COTRANSLATIONAL_PROTEIN_TARGETING_TO_MEMBRANE
GO_CYTOSOLIC_RIBOSOME
TUN - retrograde vesicle transport golgi
GO_GOLGI_VESICLE_TRANSPORT
GO_RETROGRADE_VESICLE_MEDIATED_TRANSPORT_GOLGI_TO_ENDOPLASMIC_RETICULUM
TUN - regulation defense interferon virus
GO_RESPONSE_TO_TYPE_I_INTERFERON
GO_DEFENSE_RESPONSE_TO_VIRUS
GO_REGULATION_OF_DEFENSE_RESPONSE_TO_VIRUS
GO_RESPONSE_TO_VIRUS
TUN - viral genome replication regulation
GO_NEGATIVE_REGULATION_OF_VIRAL_GENOME_REPLICATION
GO_NEGATIVE_REGULATION_OF_MULTI_ORGANISM_PROCESS
GO_REGULATION_OF_VIRAL_GENOME_REPLICATION
GO_NEGATIVE_REGULATION_OF_VIRAL_LIFE_CYCLE
GO_REGULATION_OF_VIRAL_LIFE_CYCLE
GO_REGULATION_OF_SYMBIOSIS_ENCOMPASSING_MUTUALISM_THROUGH_PARASITISM
GO_NEGATIVE_REGULATION_OF_VIRAL_PROCESS
GO_VIRAL_GENOME_REPLICATION
GO_VIRAL_LIFE_CYCLE
GO_POSITIVE_REGULATION_OF_VIRAL_LIFE_CYCLE
TUN - unfolded protein folding binding
GO_UNFOLDED_PROTEIN_BINDING
GO_PROTEIN_FOLDING

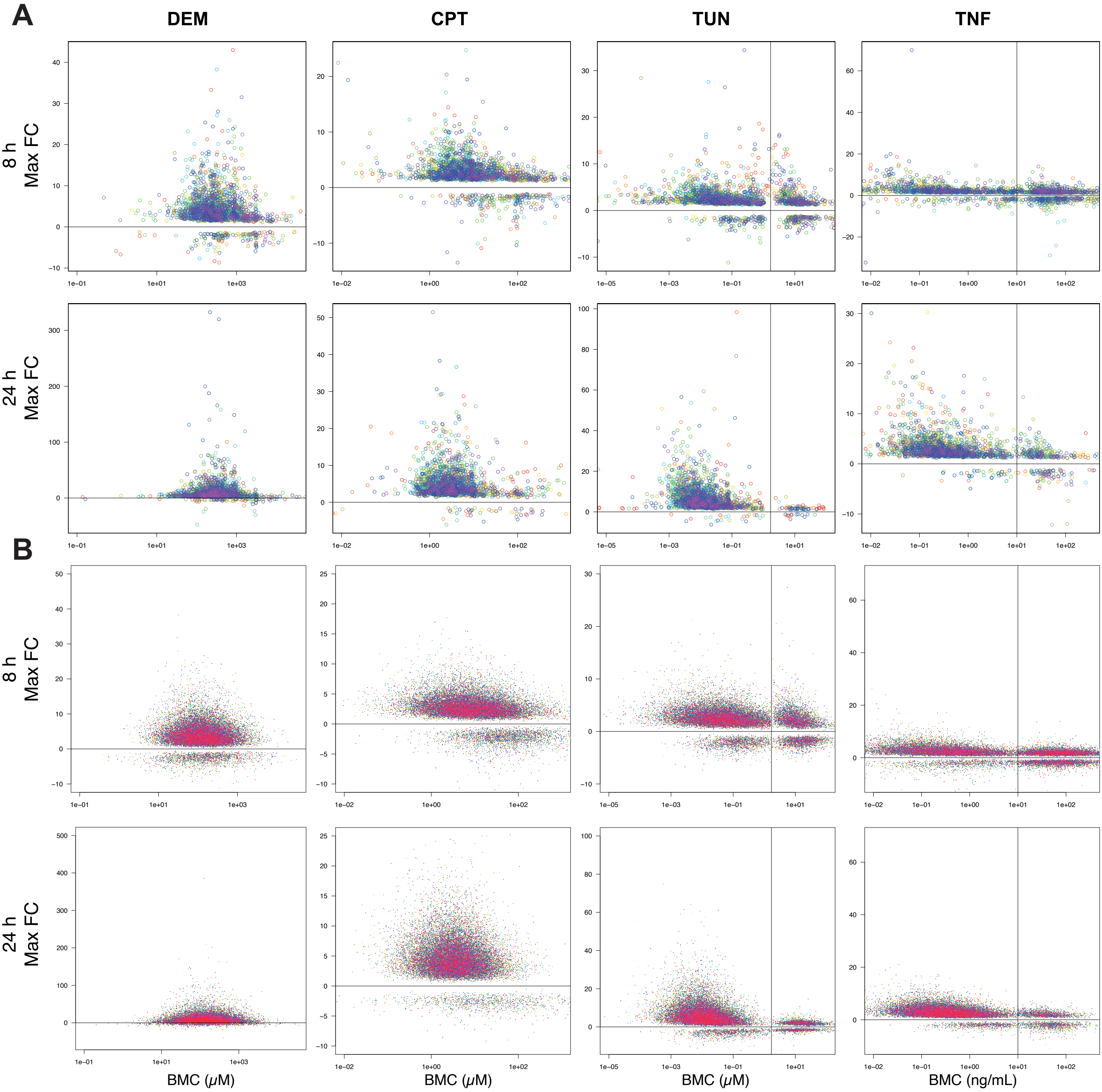
A
DEM
CPT
TUN
TNF
8 h
Max FC
24 h
Max FC
B
8 h
Max FC
24 h
Max FC
BMC (μM)
BMC (μM)
BMC (μM)
BMC (ng/mL)

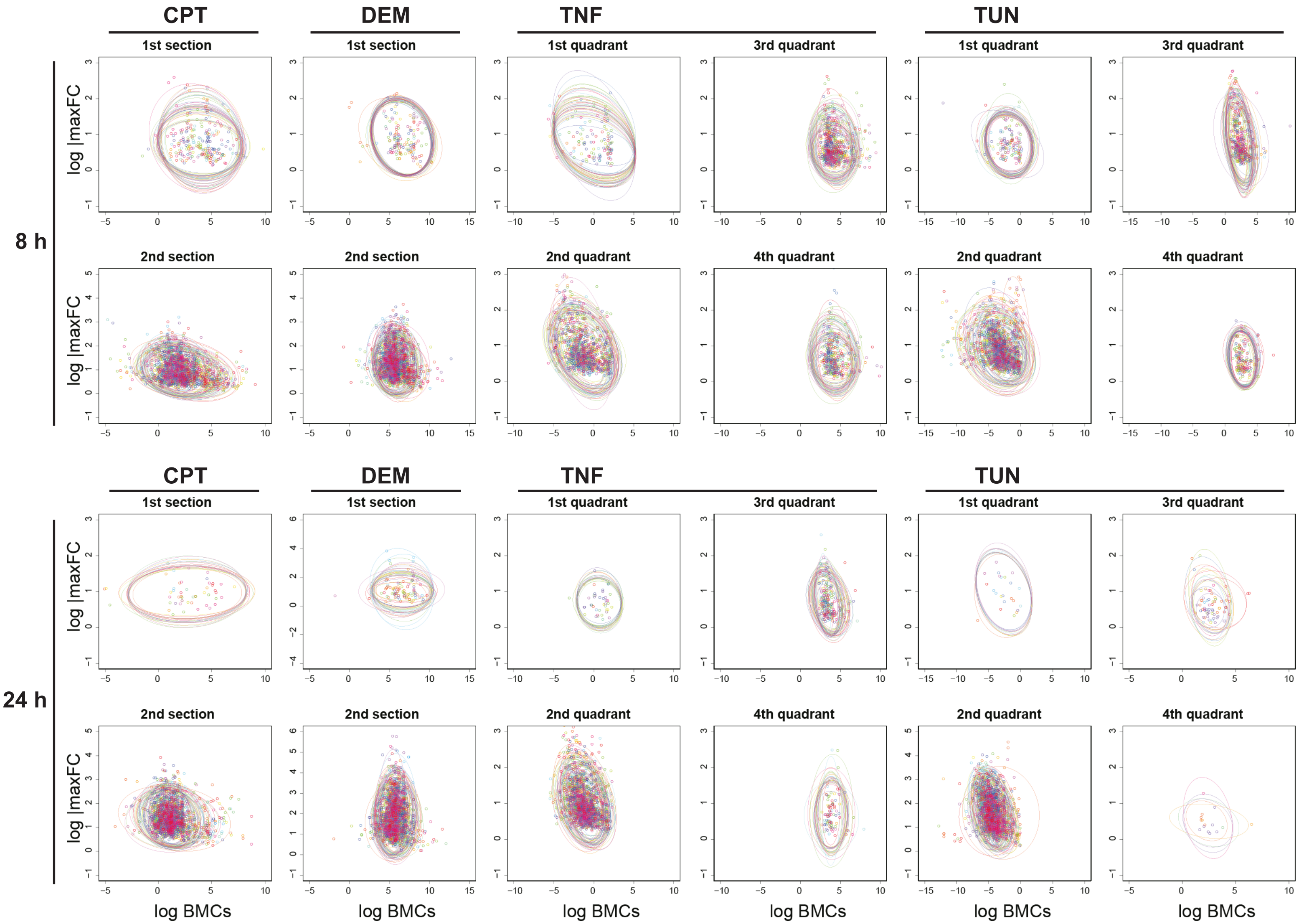
8 h
24 h
CPT
DEM
TNF
TUN
1st section
2nd section
1st quadrant
2nd quadrant
3rd quadrant
4th quadrant
log |maxFC
log BMCs
